# Truncation of poly-*N*-acetylglucosamine (PNAG) polymerization with *N*-acetylglucosamine analogues

**DOI:** 10.1101/2021.10.13.464265

**Authors:** Zachary A. Morrison, Alexander Eddenden, Adithya Shankara Subramanian, P. Lynne Howell, Mark Nitz

## Abstract

Bacteria require polysaccharides for structure, survival, and virulence. Despite the central role these structures play in microbiology few tools are available to manipulate their production. In *E. coli* the glycosyltransferase complex PgaCD produces poly-*N*-acetylglucosamine (PNAG), an extracellular matrix polysaccharide required for biofilm formation. We report that C6-substituted (H, F, N_3_, SH, NH_2_) UDP-GlcNAc substrate analogues are inhibitors of PgaCD. *In vitro* the inhibitors cause PNAG chain termination; consistent with the mechanism of PNAG polymerization from the non-reducing terminus. *In vivo*, expression of the GlcNAc-1-kinase NahK in *E. coli* provided a non-native GlcNAc salvage pathway that produced the UDP-GlcNAc analogue inhibitors *in situ*. The 6-fluoro and 6-deoxy derivatives were potent inhibitors of biofilm formation in the transformed strain, providing a tool to manipulate this key exopolysaccharide. Characterization of the UDP-GlcNAc pool and quantification of PNAG generation support PNAG termination as the primary *in vivo* mechanism of biofilm inhibition by 6-fluoro UDP-GlcNAc.

## Introduction

For bacteria, glycans form the skeleton, and the outermost contacts with their environment making these structures key to survival, and virulence.^1–3^ The use of modified monosaccharides as tools to study the biochemistry of bacterial glycans is evolving but lags behind the sophistication of experiments carried out in eukaryotic systems.^4,5^ This is in large part due to the diversity of glycan structures in bacteria and the differences in bacterial monosaccharide metabolic pathways.^6^ The incorporation of modified bacterial sugars such as 3-deoxy-D-manno-octulosonic acid (KDO),^7^ pseudaminic acid,^8,9^ legionaminic acid and trehalose^10–12^, which are salvaged and presented by bacteria in specific glycans, have provided innovative bacterial imaging methods. Similarly, other modified monosaccharides are taken up and incorporated into glycans in a species-specific manner, stimulating research into the metabolic pathways and structures that are produced.^13–18^ Engineering of bacteria to introduce novel monosaccharide salvage pathways has led to the production of tagged fucosylated O-antigens and modified peptidoglycan structures.^19–21^ Recently, the metabolic engineering of bacteria combined with a modified muramic acid residue has demonstrated the potential of monosaccharide tools in monitoring peptidoglycan biosynthesis in *H. pylori*.^22^ Here we demonstrate that metabolic engineering of the *E. coli* GlcNAc salvage pathway and the use of GlcNAc derivatives provides a method to control the synthesis of poly-*N*-acetylglucoasamine (PNAG), a key bacterial exopolysaccharide in biofilm formation.^23–32^

The generation of biofilms contributes to the virulence of bacterial pathogens, providing resistance to the host immune response and antibiotics.^33–35^ In *E. coli* the cyclic di-GMP dependent glycosyltransferase complex PgaCD polymerizes uridine 5′-diphosphate-GlcNAc (UDP-GlcNAc) into PNAG at the inner membrane.^36,37^ However, despite its importance in bacterial virulence, the direction of polymer elongation has not been defined. PgaC has moderate homology with *E. coli* cellulose synthase BcsA (24% identity, 41% similarity) and *Rhizobium leguminosarum* chitin synthase NodC (25% identity, 45% similarity), which both elongate their polysaccharides from the non-reducing end.^38–41^ However, it also has homology with *Streptococcus pyogenes* hyaluronan synthase HAS (23% identity, 45% similarity) which elongates from the reducing end.^42^ Using UDP-GlcNAc derivatives as inhibitors of PNAG biosynthesis the direction of polysaccharide elongation can be established.

The inhibition, and the mechanisms of glycosyltransferases have been investigated by generating glycosyl acceptors where the key hydroxyl has been replaced by alternative functionalities such as deoxy, fluoro or amino substituents.^43–45^ The earliest examples of this approach demonstrated deoxy acceptor analogues acting as competitive inhibitors with K_i_ values close to the K_m_ of the respective substrates.^43^ In some cases, amino substituted acceptor analogues were more potent inhibitors but had complex inhibition profiles.^45,46^ More recently, in the case of polymerizing glycosyltransferases, C5 and C6 deoxy UDP-galactofuranose analogues were shown to be potent inhibitors of mycobacterial galactan biosynthesis, and fluorinated analogues at C5 or C6 allowed for the specificity of glycosyl linkage formation by GlfT2 to be investigated.^47,48^ In both cases, the modified UDP-galactofuranose served as a substrate, that was added to the galactan chain. This modified acceptor prevented subsequent elongation and truncated polymerization. Similarly, chain termination of *Pastuerella multocida* heparosan synthase 1 with UDP-4-fluoroGalNAc has been reported.^49^ The challenges involved in delivering the donor analogue has prevented most of these approaches from being translated into *in vivo* tools, but successes have been reported. For example, delivery of an amino functionalized acceptor analogue has been shown to block the biosynthesis of the blood group A determinant^50^ and GPI anchor biosynthesis can be truncated in *Trypanosoma brucei* and *Plasmodium falciparum* with 2-amino-2-deoxy-mannose, albeit at high inhibitor concentrations.^51,52^

Herein, we have synthesized UDP-GlcNAc substrate analogues in which the C6 hydroxyl acceptor has been substituted with hydrogen, thiol, fluorine, amine or azide and demonstrate that they are PgaCD inhibitors (**2-X, Fig. 1**). The deoxy and fluoro derivatives served as transferase substrates forming terminated polymers which are potent inhibitors of PgaCD. This inhibition mechanism provides direct support for chain elongation of PNAG at the non-reducing terminus. Using an engineering approach, an unnatural GlcNAc salvage pathway was introduced in *E. coli*, allowing the C6 modified GlcNAc derivatives (**1-X, Fig. 1**) to be used *in vivo* at low micromolar concentrations for inhibition of PNAG synthesis and biofilm formation.

**Figure 1.**
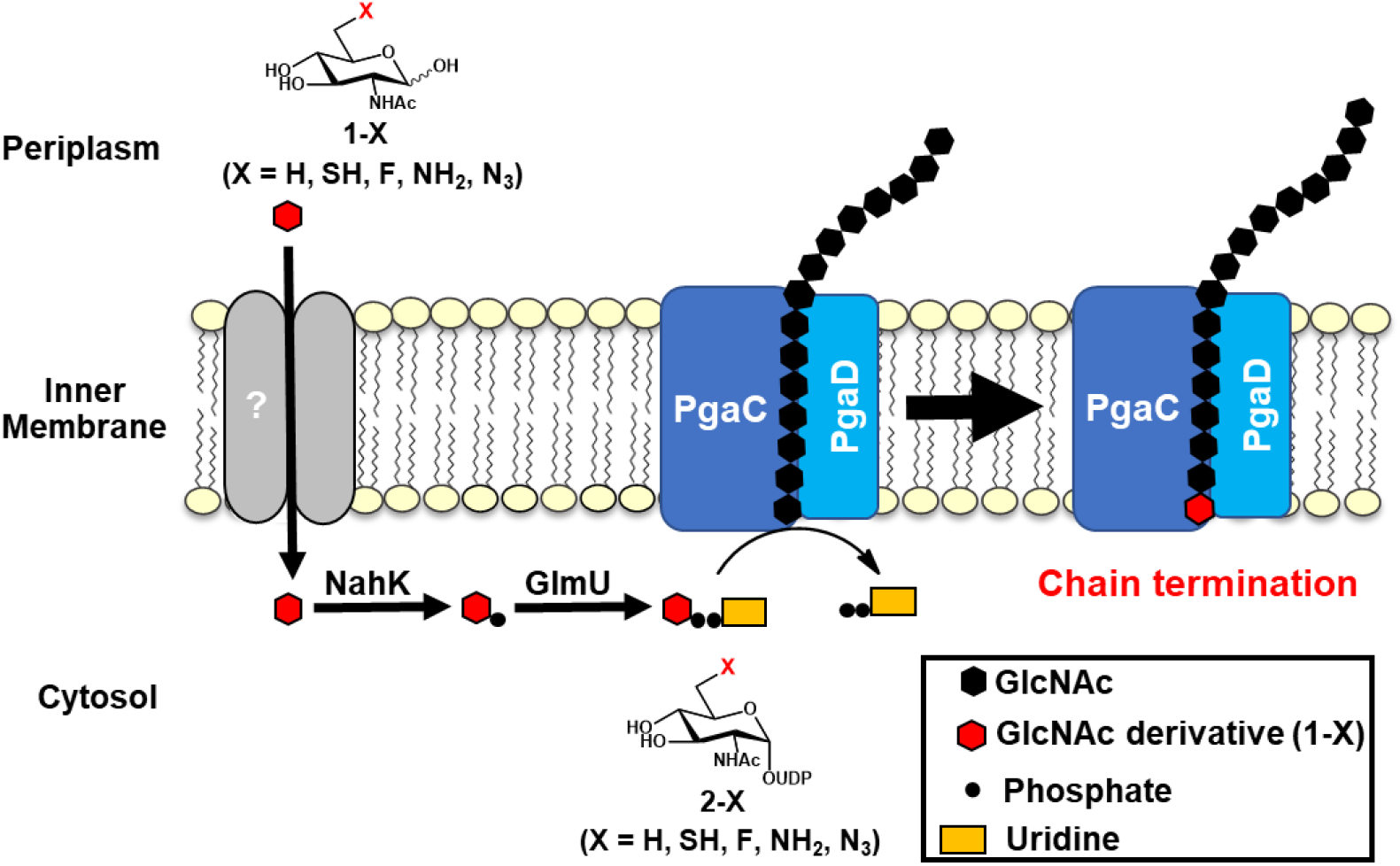
Inhibition of PNAG synthesis using C6-substituted GlcNAc analogues. Analogues **1-X** access the *E. coli* cytoplasm and are converted to UDP donors (**2-X**) through an engineered salvage pathway. The mechanism of transport of **1-X** is unknown. As PgaCD glycosyl donor substrates these derivatives serve as chain terminators of PNAG polymerization.

## Results

### Assay development and purification of PgaCD

The kinetics and bis-(3’-5’)-cyclic dimeric guanosine monophosphate (c-di-GMP) dependence of PgaCD have previously been studied using crude *E. coli* membrane preparations.^37^ We observed complicating background hydrolysis of UDP-GlcNAc when using these preparations. To overcome this challenge, we recombinantly overexpressed and purified PgaCD from *E. coli* BL21 cells. Our initial attempts to purify the wild-type PgaCD complex were unsuccessful due to protein aggregation. We reasoned that the instability and aggregation of the complex may be due to the absence of c-di-GMP in the purification buffers as complex formation of the wild-type protein has been demonstrated to be dependent on this ligand.^37^ As a single point mutation in PgaC (V227L) has been shown to partially uncouple PgaCD transferase activity from c-di-GMP and the double PgaD (N75D, K76E) mutant fully releases the PgaCD complex from its c-di-GMP dependency,^37^ we next attempted to purify PgaC^V227L^PgaD^N75D/K76E^, referred to herein as PgaCD_m_. Overexpression of PgaCD_m_ and nickel affinity chromatography enabled coelution of PgaC and PgaD, and the purification of the PgaCD_m_ complex (**Fig. S1A**). This preparation of PgaCD_m_ was used for our initial IC_50_ determinations with the UDP-GlcNAc derivatives (**2-X**).

To follow the activity of our purified preparations we developed a dot blot assay using commercial wheat germ agglutinin (WGA)-horse radish peroxidase (HRP) conjugates. WGA has affinity for GlcNAc-containing glycans and thus is expected to bind PNAG polymers. Incubation of membrane preparation or purified PgaCD_m_ with UDP-GlcNAc led to a strong WGA signal that was absent in our negative controls (**Fig. S1B-C**). The specificity of the assay was further probed using short synthetic oligomers^53^ and Dispersin B (DspB), a PNAG specific hexosaminidase which exclusively hydrolyzes GlcNAc-β-1,6 linkages.^54^ Using synthetic oligomers we determined that oligosaccharides of PNAG up to 10 GlcNAc units could not be robustly detected with WGA-HRP, likely due to limited binding of the oligosaccharides to nitrocellulose (**Fig. S1D-E**). Co-incubation of the reaction mixture with DspB completely eliminated the WGA-HRP signal. This suggests that that the PNAG produced by membrane preparations of PgaCD_m_ is significantly longer than 10 residues. DspB has been shown to disrupt PNAG dependent biofilms by cleaving PNAG into monomers and shorter oligomers.^55^

### C6-substituted UDP-GlcNAc derivatives inhibit PgaCD in vitro

Given the previous successes of modified glycosyl acceptors as inhibitors of glycosyl transferases^43–52^ and the possible utility of these compounds in determining the elongation direction of PgaCD, a chemoenzymatic route was used to produce the series of C6 functionalized UDP-GlcNAc derivatives (**Fig. 1, 2-X**).^56^ Using the WGA dot blot assay, quantified by densitometry, the inhibitory activity of the substrate analogues was evaluated against purified PgaCD_m_ **(Fig. S2**). More conservative C6 substitutions gave greater inhibition against PNAG production: 6-deoxy and 6-thio derivatives were most potent, followed by the 6-fluoro and lastly the 6-amino derivative (**Fig. 2**). The 6-azido analogue was inactive up to 10 mM, suggesting that PgaCD’s active site cannot accommodate excessive steric bulk around the C6 acceptor. Interestingly, all active UDP-GlcNAc derivatives were substantially more potent inhibitors than UDP which is often found to display product inhibition in glycosyltransferase assays.

**Figure 2.**
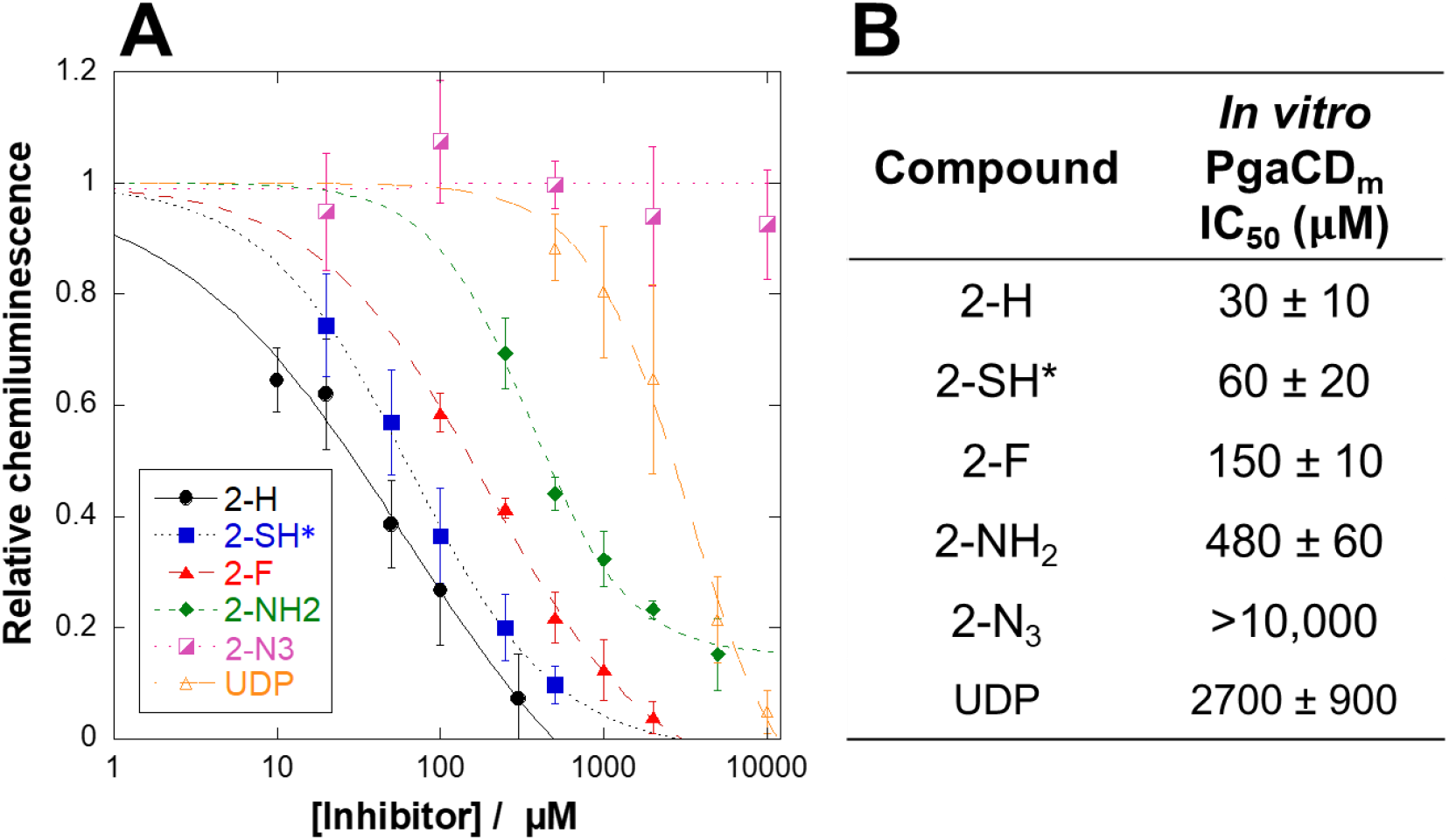
C6-substituted UDP-GlcNAc derivatives inhibit PgaCD_m_. **A)** Dose-response curves for the inhibition of purified PgaCD_m_ by C6-substituted UDP-GlcNAc analogues. **B)** IC_50_ values measured by dot blot assay (*n = 3)*. The uncertainties are standard deviations between replicates. *Assayed in presence of 5 mM DTT.

### Mechanism of UDP-6-deoxyGlcNAc inhibition and polymerase elongation

Depending on the elongation direction of PgaCD catalyzed PNAG polymerization, we hypothesized the most potent inhibitor UDP-6-deoxyGlcNAc may act either as a chain terminator or as a competitive inhibitor of PNAG elongation. Kinetic experiments were carried out to discriminate between these modes of inhibition. Chain termination is only possible with non-reducing end PNAG elongation since the analogues lack a C6 hydroxyl acceptor. Given the long chain length of PNAG produced *in vivo*, PgaCD is expected to have high processivity.^57,58^ Thus, release of the chain terminated PNAG product may be slow, leading to the observation of irreversible inhibition.

Initial experiments with purified PgaCD_m_, in comparison to the crude PgaCD_m_ membrane preparation (MP), showed significantly more polymerase activity in the membrane preparation (**Fig. S1C**) and that the stability of the activity over the course of the reaction was superior in the membrane preparation. Thus, for the time course experiments the more active membrane preparation PgaCD_m_(MP) was used.^37^ We found that our WGA-HRP dot blot assay was suitable for monitoring the rate of PNAG synthesis over extended incubation periods (24 h).

Time dependent inhibition of PgaCD_m_(MP) was observed with UDP-6-deoxyGlcNAc. As the reaction progressed product formation was observed to plateau prior to reaction completion (**Fig. 3A**). This observation is incompatible with a competitive inhibitor which would be expected to slow the catalysis consistently over the time course. Furthermore, the inhibitor’s high potency in this experiment despite the large excess of UDP-GlcNAc (10 mM) also argued against competitive inhibition. For instance, at 50 μM inhibitor and 10 mM UDP-GlcNAc >50% inhibition of PNAG production was observed. A similar level of inhibition was measured in the IC_50_ experiments using only 300 μM UDP-GlcNAc (IC_50_ = 30 μM, **Fig. 2B**). It was not possible to rule out loss of enzyme activity and substrate depletion due to other enzyme activities in the preparation contributing to the observed plateaus in product formation, but the linear increase in PNAG formation observed over 8 h in the absence of inhibitor suggests these contributions were minimal under the experimental conditions.

**Figure 3.**
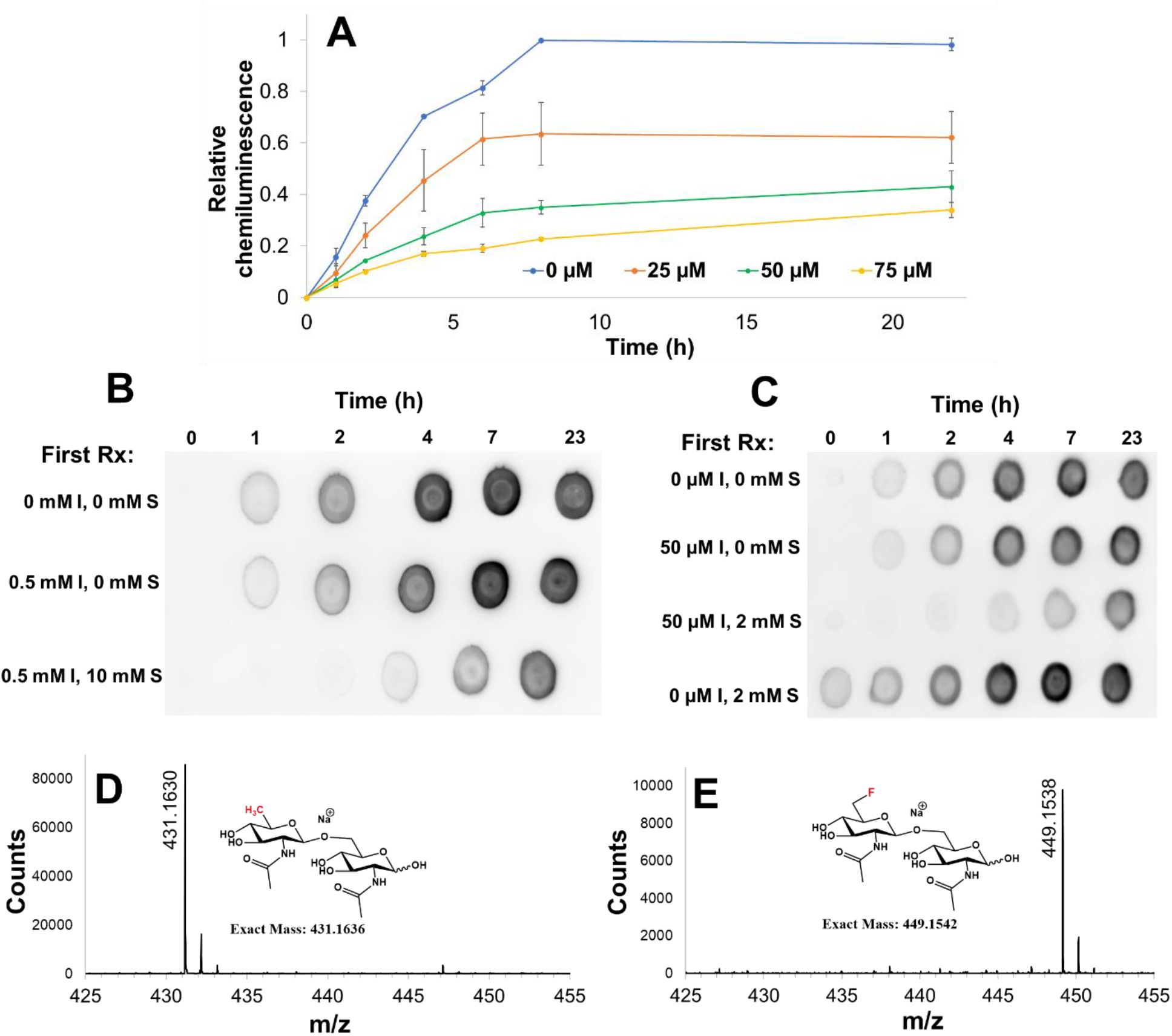
UDP-6-deoxyGlcNAc inhibits PgaCD through chain termination. **A**) Time course of PgaCD_m_(MP) reactions in the presence of UDP-GlcNAc (10 mM) and varying amounts of UDP-6-deoxyGlcNAc (0 – 75 μM). The values are the averages of two biological replicates, the error bars are standard deviations. **B**) Overnight incubation of PgaCD_m_(MP) with a mixture of UDP-6-deoxyGlcNAc (I) and UDP-GlcNAc (S) causes inhibition that is not overcome by dialysis. Post-dialysis reaction conditions: 10 mM UDP-GlcNAc, 5 mM MgCl_2_, 37 °C. **C**) Overnight incubation of PgaCD_m_(MP) with a mixture of UDP-6-deoxy-GlcNAc (I) and UDP-GlcNAc (S) causes inhibition that is not overcome by 10-fold dilution. Post-dilution reaction conditions: UDP-GlcNAc (20 mM), MgCl_2_ (5 mM), 37 °C. Biological replicate of the experiments in panels B and C gave qualitatively similar results. **D-E**) ESI-HRMS confirms formation of expected disaccharide products from reactions of incubating GlcNAc (50 mM) and UDP-6-deoxyGlcNAc (5 mM) (**D**) or UDP-6-fluoroGlcNAc (5 mM) (**E**) with PgaCD_m_(MP). The samples were purified by Carbograph column prior to mass spectrometry analysis.

Evidence of time dependent irreversible inhibition was also observed in a dialysis experiment. The PgaCD_m_(MP) was incubated overnight with or without UDP-6-deoxyGlcNAc and UDP-GlcNAc. The samples were dialyzed to remove the inhibitor and then fresh excess UDP-GlcNAc was added and the reaction was monitored by dot blot assay (**Fig. 3B**). When the enzyme was preincubated with inhibitor alone, no appreciable inhibition of PNAG production resulted after dialysis. However, when the preincubation contained both inhibitor and substrate a dramatic reduction in PNAG production was observed after dialysis. The requirement for UDP-GlcNAc and the inability of dialysis to restore activity suggests a chain termination mechanism followed by slow release of the truncated polymer.

A dilution experiment was performed to provide further support for the chain termination mechanism. The PgaCD_m_(MP) was incubated overnight with or without UDP-GlcNAc and UDP-6-deoxyGlcNAc. The reactions were diluted 10-fold, to a concentration where the inhibitor has minimal activity, additional UDP-GlcNAc was added, and PNAG production was monitored (**Fig. 3C**). Consistent with the dialysis experiment, incubating PgaCD_m_(MP) with inhibitor in the absence of UDP-GlcNAc gave no appreciable reduction in PNAG formation, confirming that the dilution was adequate to prevent inhibition. A reduction in PNAG production was observed only when the enzyme was incubated with both inhibitor and substrate, further supporting formation of an inactive PgaCD_m_/chain-terminated PNAG complex.

Taken together, a competitive inhibition model cannot account for these observations: the inhibition requires the presence of substrate and has irreversible character. We conclude that UDP-6-deoxyGlcNAc inhibits PgaCD through chain termination, requiring that PgaCD elongates PNAG from its non-reducing end.

The proposed inhibition mechanism requires that PgaCD utilizes the C6-substituted UDP-GlcNAc analogues as glycosyl donors truncating the nascent PNAG chain. However, chain terminated PNAG oligomers could not be detected by mass spectrometry or ^19^F NMR experiments, likely because of slow release of the truncated oligomers from the enzyme limiting product accumulation. We hypothesized that if GlcNAc were used as an acceptor in the reaction with the C6 donor analogue the disaccharide product would be sufficiently small to dissociate from PgaCD_m_, permitting multiple catalytic cycles and accumulation of the disaccharide in the reaction mixture. Incubating GlcNAc (50 mM) and UDP-6-deoxyGlcNAc or UDP-6-fluoroGlcNAc (5 mM) with PgaCD_m_(MP) gave product masses consistent with the expected disaccharide products (m/z = 431.1630 or 449.1538) further supporting the chain termination mechanism and that PNAG elongation occurs from the non-reducing terminus (**Fig. 3D-E**).

### C6-substituted UDP-GlcNAc derivatives inhibit PNAG production and biofilm formation

We hypothesized that these UDP-GlcNAc derivatives may be active against PgaCD *in vivo*. Recognizing that direct cellular entry of nucleotide sugars was unlikely, we envisioned feeding precursors which could generate the desired inhibitor *in situ. E. coli* is not known to have GlcNAc-1-kinase activity and GlcNAc salvage proceeds via phosphorylation at C6, deacetylation by NagA, and conversion to GlcN-1P by GlmM, prior to reacetylation and uridyl transfer by GlmU.^59^ The C6-substituted GlcNAc derivatives are not expected to transverse this pathway. To bypass these metabolic steps, we reasoned that we could obtain the desired UDP-GlcNAc derivatives in the bacterial cytoplasm if a novel salvage pathway for these derivatives could be introduced.

NahK is a promiscuous GlcNAc-1-kinase that has been used extensively for the chemoenzymatic synthesis of GlcNAc-1P derivatives.^56,60–62^ Through the introduction of NahK activity to the bacteria we envisaged that the C6 GlcNAc derivatives **1-X** could be phosphorylated providing an unnatural salvage pathway to the desired inhibitors **2-X** *in situ* (**Fig. 1**). NahK substrate specificity has been thoroughly studied and C6-subtituted GlcNAc analogues are known to be phosphorylated.^56^ The resulting 1-phosphates are then expected to be converted to their UDP donors by *E. coli* GlmU.^56^ To evaluate this strategy we used *E. coli* MG1655 *csrA*::*kanB* as a model system as this strain is known to constitutively form a PNAG-based biofilm.^36,63^

For control of NahK activity, an L-arabinose-inducible pBAD plasmid encoding recombinant His-tagged NahK was introduced into *E. coli csrA*::*kanB*. A plasmid encoding a catalytically inactive NahK^D208N/K210E/N213A^ triple mutant, referred to herein as NahK_m_, was also developed as a negative control for GlcNAc-1-kinase activity. Induction with L-arabinose in plasmid-containing *E. coli csrA*::*kanB* gave a high level of NahK or NahK_m_ expression and there was minimum expression in its absence (**Fig. S3**). Induction of either protein had no appreciable effect on biofilm formation relative to the parent strain *E. coli csrA::kanB*, and cells expressing NahK or NahK_m_ had equivalent growth curves (**Fig. S4**).

Feeding all C6-substituted GlcNAc derivatives led to NahK-dependent biofilm inhibition (**Fig. 4A-C** & **Table 1**). When NahK was not induced or when inactive NahK_m_ was expressed minimum inhibition was observed suggesting that 1-phosphorylation of the GlcNAc derivatives is required for their activity and minimal effects arise from NahK expression. The most active compounds were 6-deoxy and 6-fluoroGlcNAc (IC_50_ ∼10 μM). Interestingly, though UDP-6-thioGlcNAc was among the most potent *in vitro* PgaCD_m_ inhibitors, this analogue was somewhat less active *in vivo* (IC_50_ ∼100 μM). This may be a consequence of disulfide formation limiting its cellular entry. In support of this, conducting the biofilm assay in the presence of DTT (0.1 mM) halved its IC_50_ value. Furthermore, 6-aminoGlcNAc had similar activity to 6-thioGlcNAc, consistent with its modest potency *in vitro*. It is notable that 6-azidoGlcNAc, inactive *in vitro*, was also least potent *in vivo*, inhibiting biofilm formation only at millimolar concentrations. This correlation supports our hypothesis that the UDP-GlcNAc derivatives (**2-X**) truncate PNAG polymerization *in vivo*.

**Table 1.**
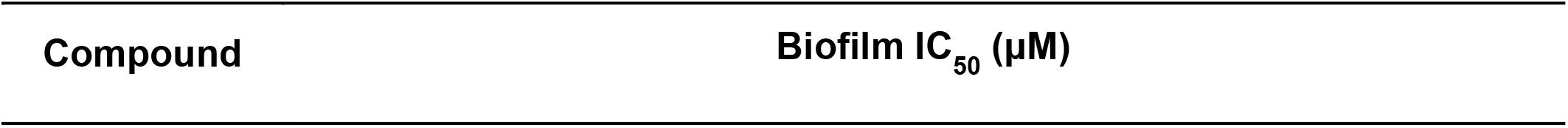
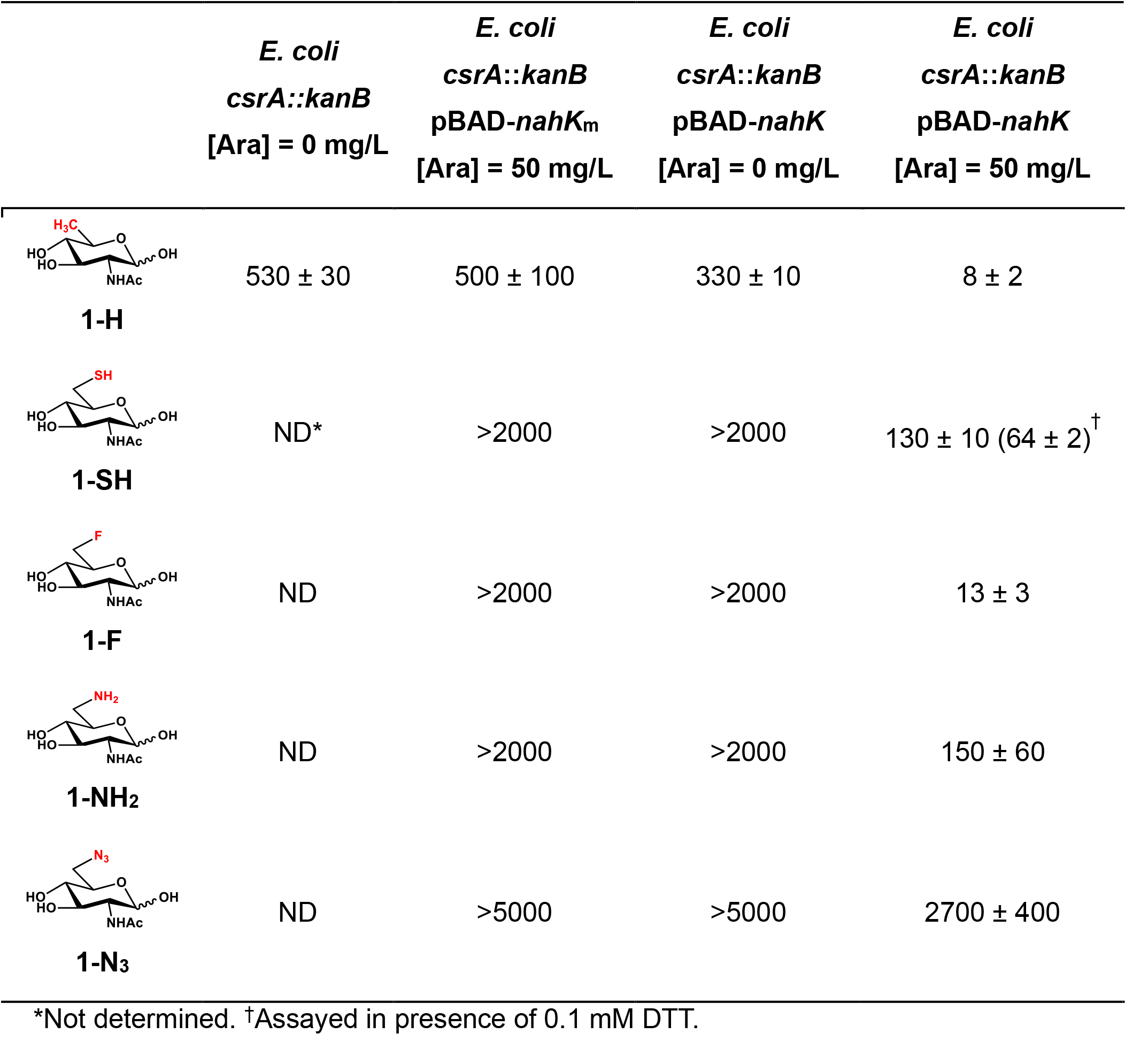
Biofilm IC_50_ values against *E. coli csrA::kanB*. The values are the averages of two biological replicates, the uncertainties are standard deviations.

|  | *E. coli*<br>*csrA::kanB*<br>[Ara] = 0 mg/L | *E. coli*<br>*csrA::kanB*<br>pBAD-*nahK<sub>m</sub>*<br>[Ara] = 50 mg/L | *E. coli*<br>*csrA::kanB*<br>pBAD-*nahK*<br>[Ara] = 0 mg/L | *E. coli*<br>*csrA::kanB*<br>pBAD-*nahK*<br>[Ara] = 50 mg/L |
| --- | --- | --- | --- | --- |
| **1-H** | 530 ± 30 | 500 ± 100 | 330 ± 10 | 8 ± 2 |
| **1-SH** | ND* | >2000 | >2000 | 130 ± 10 (64 ± 2)<sup>†</sup> |
| **1-F** | ND | >2000 | >2000 | 13 ± 3 |
| **1-NH<sub>2</sub>** | ND | >2000 | >2000 | 150 ± 60 |
| **1-N<sub>3</sub>** | ND | >5000 | >5000 | 2700 ± 400 |

\*Not determined. <sup>†</sup>Assayed in presence of 0.1 mM DTT.

**Figure 4.**
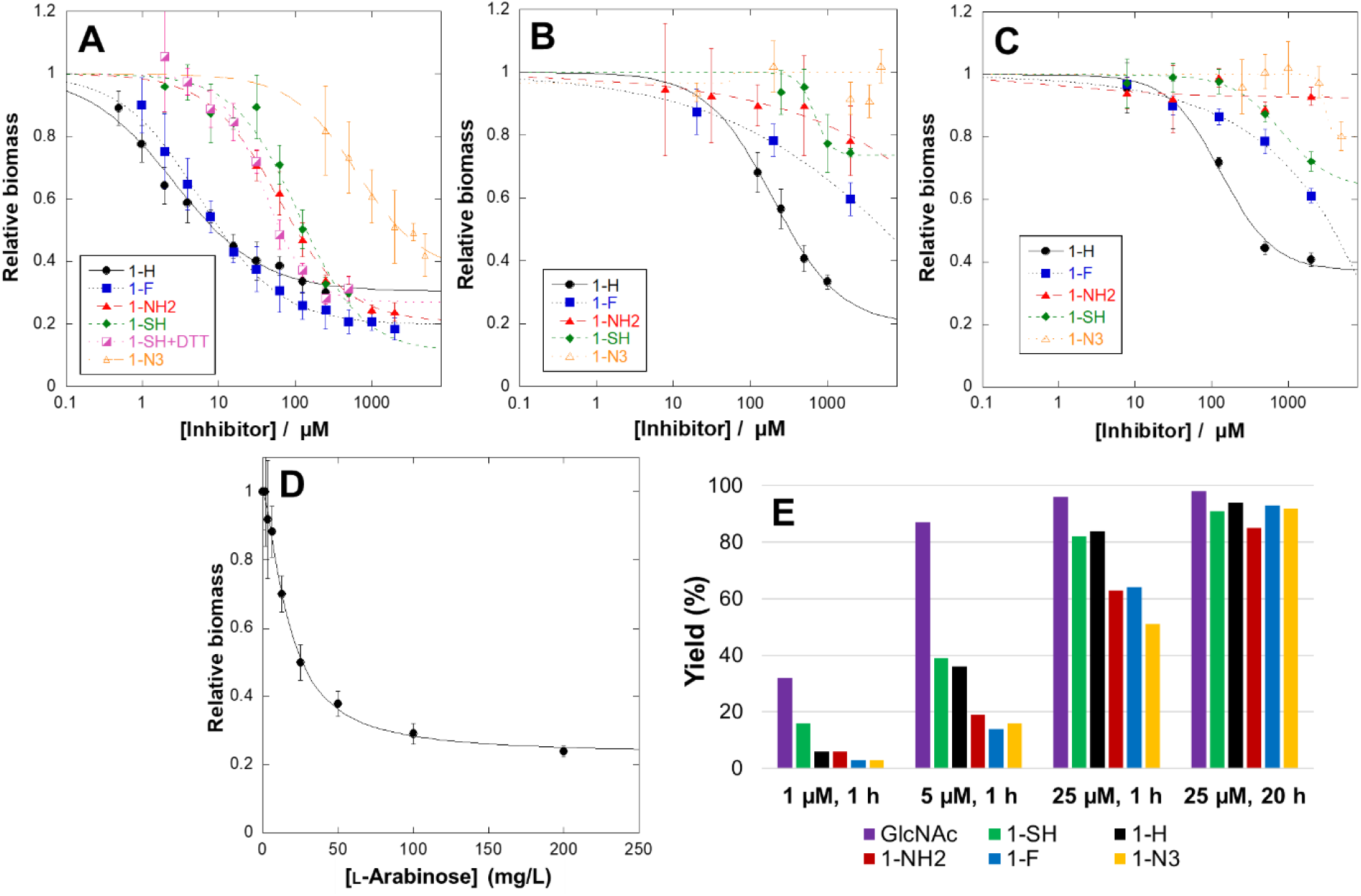
NahK activity potentiates biofilm inhibition by C6-substituted GlcNAc derivatives. **A**) Dose-response curves for biofilm inhibition of *E. coli csrA*::*kanB* pBAD-*nahK* induced with arabinose by C6-substituted GlcNAc derivatives. Limited inhibition was seen with uninduced pBAD-*nahK* (**B**) or induced inactive pBAD-*nahK*_m_ (**C**). The values are the averages of 4-6 technical replicates and the uncertainties are standard deviations. Inductions were done at 50 mg/L L-arabinose. **D**) Dependence of 6-fluoroGlcNAc (20 μM) biofilm inhibition on L-arabinose concentration. The values are the average of four technical replicates and the error bars are standard deviations. **E**) Relative NahK (1, 5, or 25 μM) phosphorylation rates of C6-substituted GlcNAc derivatives (10 mM) estimated by ^1^H NMR assay.

### In vivo mechanism of action studies

To clarify the *in vivo* mechanism of action of these GlcNAc derivatives, we first studied their effect on growth rate (**Fig. S5**). The 6-deoxy, 6-fluoro, 6-amino, and 6-thio derivatives were found to have minimum effect on *E. coli* growth at the concentrations required to inhibit biofilm formation. Given the limited toxicity observed it is unlikely that these derivatives are being incorporated in appreciable quantities in LPS as 1→6 linkages are key to the core disaccharide and LPS is essential for growth.^3^ It is possible that the compounds are being incorporated into peptidoglycan, but it appears to be non-perturbing to growth and given the importance of 6-OH in lytic transglycosylase activity as well as peptidoglycan recycling significant incorporation is unlikely.^65^ Interestingly, 5 mM 6-azidoGlcNAc caused *E. coli* growth inhibition, but only during mid to late log phase. This observation helps account for the biofilm inhibition at these high concentrations despite the inactivity of its UDP derivative against PgaCD_m_ *in vitro*.

Though both 6-fluoro and 6-deoxyGlcNAc had promising potencies against biofilm formation, the activity of the 6-fluoro derivative gave a greater change in biofilm inhibition upon NahK induction, showing no inhibition at 2 mM in the absence of NahK. Therefore, we sought to further characterize the *in vivo* mechanism of 6-fluoroGlcNAc.

We first examined the dependence of 6-fluoroGlcNAc biofilm inhibition on the NahK expression levels by varying the arabinose concentration in the presence of 6-fluoroGlcNAc (20 μM) (**Fig. 4D**). The inhibition was highly dependent on the NahK induction level, plateauing only at arabinose concentrations above ∼50 mg/L (EC_50_ = 15 ± 3 mg/L). An *in vitro* ^1^H NMR assay was subsequently used to determine the relative rates of phosphorylation of the C6-substituted GlcNAc derivatives by NahK (**Fig. 4E**). We found that 6-fluoroGlcNAc was one of the least favourable substrates, as can be seen by comparing the product formation over time with increasing enzyme concentrations. At long time points with high enzyme concentrations the desired UDP-GlcNAc derivatives are produced in all cases and we find that UDP-6F-GlcNAc is produced *in vivo* (see below). This suggests that at the chosen 50 mg/L arabinose concentration significant phosphorylation of all the GlcNAc derivatives (**1-X**) is likely occurring *in situ*.

Next the generation of the respective UDP-GlcNAc derivative *in situ* was investigated. In conjunction with the generation of the UDP-GlcNAc derivative we were curious if the biosynthesis and accumulation of unnatural UDP-GlcNAc analogues would adversely affect the size of the bacteria’s UDP-GlcNAc pool. Notably, the reported K_m_ value of PgaCD for UDP-GlcNAc (∼0.3 mM)^37^ is larger than typical estimates of its pool level in *E. coli* (∼0.1 mM),^66^ implying PNAG biosynthesis would be sensitive to changes in the UDP-GlcNAc level. There is precedence for this type of inhibition mechanism, UDP-6-fluoroGalNAc was found to reduce glycosaminoglycan biosynthesis in eukaryotes primarily through depletion of the UDP-GalNAc pool.^67^ Similarly, 3-F sialic acid perturbed the concentrations of CMP-Sia in cells leading to the loss of cell surface sialic acid.^68^ To quantify the level of UDP-GlcNAc in *E. coli csrA*::*kanB* pBAD-*nahK* we adapted an established reverse phase ion-pairing HPLC method using potassium phosphate buffer with tetrabutylammonium phosphate.^69,70^ Though two columns in series were required in these previous reports, we found that adequate resolution of the UDP-GlcNAc peak could be achieved by using a single Restek Ultra IBD column and optimizing the mobile phase potassium phosphate concentration. Standard addition of UDP-GlcNAc to an *E. coli csrA*::*kanB* pBAD-*nahK* metabolite extract confirmed the identity and homogeneity of the assigned peak (**Fig. S6A**).

Using this HPLC method, we evaluated the production of UDP-6-fluoroGlcNAc (0, 20, or 200 μM) and the concentration of UDP-GlcNAc in the presence of arabinose (50 mg/L) in the *E. coli csrA::kanB* pBAD-*nahK* strain (**Fig. 4, S6B & S7**). At 20 μM there was no change in the UDP-GlcNAc level and only small amounts of UDP-6-fluoroGlcNAc were detected. However, at 200 μM a decrease to ∼68% of the untreated lysate UDP-GlcNAc concentration was observed and the UDP-6-fluoroGlcNAc concentration was similar to that of UDP-GlcNAc. From these results we conclude that the UDP-GlcNAc pool depletion is not the cause of biofilm inhibition at low inhibitor concentrations (around IC_50_). However, this mechanism may become relevant at high inhibitor concentrations.

Finally, we confirmed the effect of 6-fluoroGlcNAc on PNAG production. We extracted PNAG from *E. coli csrA*::*kanB* pBAD-*nahK* biofilms using an established procedure^71,72^ and assessed its relative level using our dot blot assay with WGA-HRP. We found that treatment with 6-fluoroGlcNAc reduced the PNAG level, but only in the presence of NahK activity (**Fig. 6A**). As a negative control, the pBAD*-nahK* plasmid was electroporated into the *E. coli* BW25113 strain, which does not constitutively produce PNAG. Induction of pBAD*-nahK* with arabinose in this genetic background did not yield a signal in our assay, confirming that PNAG rather than some other GlcNAc-containing bacterial component was being detected in the crude *E. coli csrA*::*kanB* pBAD-*nahK* extracts. Furthermore, varying the concentration of 6-fluoroGlcNAc gave dose-dependent reduction in the HRP signal (**Fig. 6B**).

**Figure 5.**
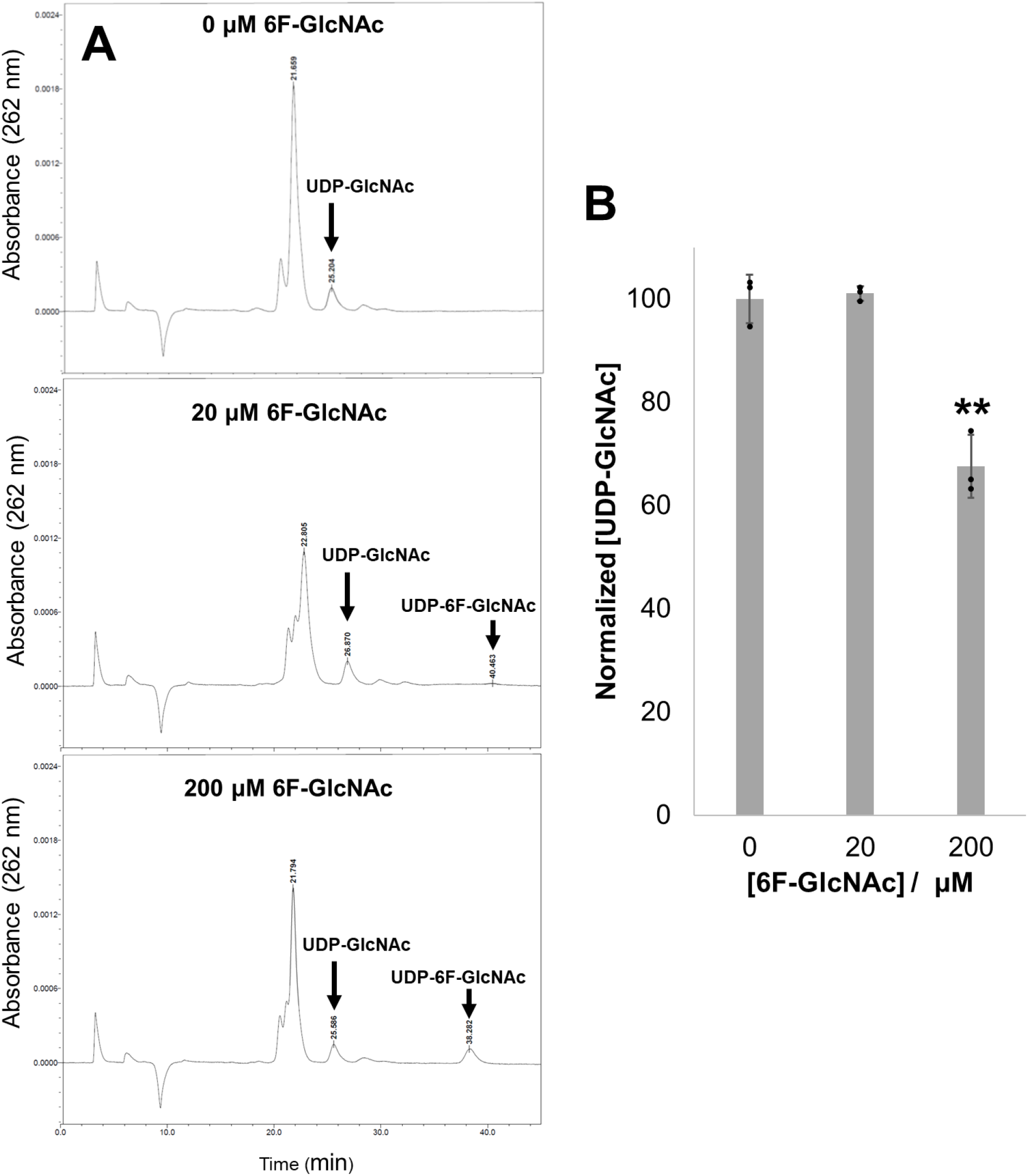
The effect of 6-fluoroGlcNAc on the UDP-GlcNAc pool in *E. coli* MG1655 *csrA::kanB* pBAD*-nahK*. **A**) Representative chromatograms of metabolite extracts prepared at 0, 20, or 200 μM 6-fluoroGlcNAc in the presence of L-arabinose (50 mg/L). Peaks were assigned using the retention times of standards ran immediately before the lysate samples. **B**) Average normalized UDP-GlcNAc levels from cells grown at different 6-fluoroGlcNAc concentrations (*n = 3*). Uncertainties are standard deviations. Statistical significance was determined using Welch’s *t*-test. **p-value < 0.01.

**Figure 6.**
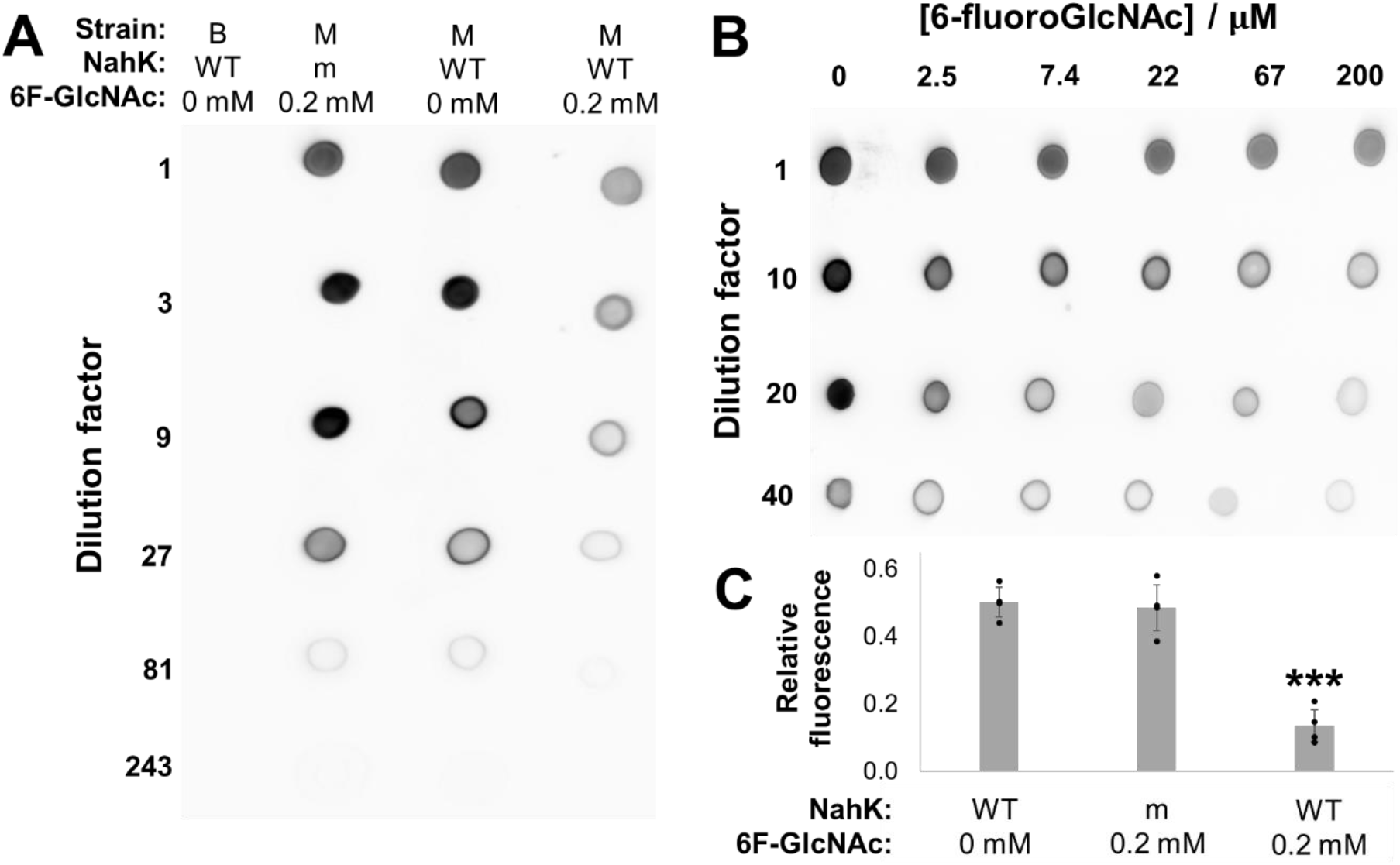
Effect of 6-fluoroGlcNAc on PNAG level. **A**) Treatment with 6-fluoroGlcNAc reduces the WGA-HRP signal of the PNAG extract only in the presence of NahK activity. Strain M refers to *E. coli* MG1655 *csrA::kanB* and strain B is *E. coli* BW25113. Additionally, WT denotes to pBAD-*nahK* and m refers to pBAD-*nahK*_m_. **B**) Dose-dependent reduction of PNAG level by 6-fluoroGlcNAc in *E. coli csrA::kanB* pBAD-*nahK* cells. pBAD-*nahK* was induced with 50 mg/L L-arabinose. Biological replicates of the experiments in panels A and B gave qualitatively identical results. **C**) 6-fluoroGlcNAc reduces GFP-DspB^E184Q^ cell surface binding exclusively in the presence of NahK activity (*n = 4*). *E. coli csrA::kanB* pBAD-*nahK* cells labelled with GFP-DspB^E184Q^ are treated with DspB to hydrolyze PNAG, and the solubilized GFP fluorescence signal is measured. pBAD-*nahK* was induced with 50 mg/L L-arabinose. Values are normalized to the fluorescence of the initial labelling solution. Uncertainties are standard deviations. Statistical significance was determined using single-factor ANOVA. ***p-value < 0.0001.

To further confirm the reduction in PNAG production we used our previously reported GFP-DspB^E184Q^ PNAG probe.^64^ PNAG-producing *E. coli csrA::kanB* pBAD-*nahK* cells were first labelled with the PNAG-binding probe, and the extent of probe release upon PNAG enzymatic hydrolysis with added DspB was used as a metric to estimate PNAG levels. Treatment with 6-fluoroGlcNAc in the presence of NahK significantly reduced the proportion of cell surface-retained GFP-DspB^E184Q^ (**Fig. 6C**), consistent with lower levels of PNAG production.

### Conclusions

Through the mechanistic investigations of C6 modified UDP-GlcNAc derivatives we have found that these compounds inhibit PNAG production by a chain termination mechanism. The terminated chains have high affinity for the PgaCD enzyme leading to a tight binding inhibitor that appears to be approaching irreversible inhibition. These results directly support that elongation of PNAG from the non-reducing terminus. This is consistent with the mechanism proposed for synthase dependent cellulose biosynthesis.^73^

Engineering a novel GlcNAc salvage pathway in *E. coli* has enabled us to use these inhibitors as tools for the control of PNAG biosynthesis *in vivo*. The introduction of the necessary GlcNAc-1P kinase activity (NahK) was minimally disruptive based on its lack of effect on biofilm formation and growth. Only in the presence of high concentrations of the GlcNAc analogue in addition to NahK were intracellular UDP-GlcNAc concentrations adversely effected. This is consistent with the known regulation of the biosynthesis of UDP-GlcNAc occurring through feedback of GlcNAc-6P and GlcN-6P.^74,75^ The mechanism that the monosaccharides use to gain access to the cytoplasm is not known and is an ongoing area of research.

Although use of these inhibitors as tools *in vivo* required the introduction of non-native NahK into the target bacteria, limiting the inhibitors therapeutic potential, the approach highlights the possible use of GlcNAc-1-phosphate derivates as antibiofilm agents when an efficient mechanism to deliver the compounds into bacteria is available. Recently, it was shown that lipid linked glycan intermediates can be directly taken up and used by mycobacteria, suggesting delivery of 1-phosphate derivatives is possible.^76^ The NahK bearing strains in combination with the inhibitors will also be useful in model studies where the temporal control of PNAG formation is required. Furthermore, beyond inhibitor delivery, the expression of NahK in *E. coli* is expected to have broad utility for achieving efficient metabolic conversion of GlcNAc derivatives. This technology may be helpful for metabolic labelling experiments and probing aspects of bacterial sugar metabolism.

## Methods

### Construction of recombinant PgaCD plasmids

The *pgaC and pgaD* genes were amplified from *E. coli K12* genomic DNA by PCR. *pgaC* and *pgaD* were cloned into pQLinkH and pQLinkN vectors, respectively, using the BamHI and NotI restriction sites. The resulting pQLinkH-6His-*pgaC* vector encodes an N-terminal hexahistidine tagged PgaC. The vector pQLinkN-*pgaD* was used as a template to introduce a 3x FLAG tag into the C-terminus of *pgaD* using round-the-horn site directed mutagenesis. Briefly, primers containing overhangs of the 3x FLAG peptide were phosphorylated using the T4 polynucleotide kinase (New England BioLabs). Phosphorylated primers were used to PCR amplify pQLinkN-*pgaD* to generate the plasmid pQLinkN-*pgaD*-FLAG. Point mutations in *pgaC* (V227L) and *pgaD* (N75D, K76E) were generated using site directed mutagenesis. The co-expression plasmid pQLink-6His-*pgaC-pgaD*-FLAG was produced by combining the two parent plasmids using ligation independent cloning.^77^ The co-expression plasmid was sequenced using internal primers for *pgaC* and *pgaD* (Centre for Applied Genomics).

### Expression and preparation of membranes overexpressing PgaCD

Chemically competent *E. coli* BL21 cells were transformed with pQLink-6His-*pgaC-pgaD*-FLAG, plated onto agar plates supplemented with 100 μg/mL carbenicillin and grown overnight at 37 °C. Transformed cells were used to inoculate 2 L of Luria broth (LB) media supplemented with 100 μg/mL ampicillin. To generate empty BL21 membranes, *E. coli* BL21 cells were plated onto agar plates and used to inoculate 2 L of LB media without antibiotic. Cells were grown at 37 °C until an optical density of 0.6-0.8 and protein expression was induced by the addition of 1 mM isopropylthio-β-galactoside (IPTG). Cultures were transferred to 18 °C and incubated overnight. Cells were harvested the following day by centrifugation at 5000 *x g* for 15 min and resuspended into 40 mL of Buffer A (50 mM HEPES pH 7.5, 300 mM NaCl, 5% v/v glycerol, 1 mM TCEP) with one protease inhibitor tablet (Sigma-Aldrich). Pellets were lysed by sonication (Misonix 3000) on an ice bath at 65% output for 1 min and 15 s. The lysate was centrifuged at 12100 *x g* for 17 min to remove cellular debris. The supernatant was further centrifuged at 257000 *x g* for 60 min at 4 °C to harvest crude membranes. The membranes were resuspended in Buffer A to a final concentration of 100 mg/mL and homogenized in a hand press. Membrane preparations were stored at −20 °C.

### Purification of PgaCD

PgaCD_m_ membranes were solubilized in Buffer A supplemented with 1% w/v DDM/0.1% w/v CHS for 60 min at 4 °C. Insoluble material was removed by centrifugation at 257000 *x g* for 60 min at 4 °C. The supernatant was applied to a gravity flow chromatography column packed with 1.5 mL of Ni Sepharose 6™ fast flow resin (GE healthcare) and incubated at 4 °C for 30 min. The resin was washed twice with 40 mL of Buffer B (50 mM HEPES pH 7.5, 300 mM NaCl, 5% v/v glycerol, 1 mM TCEP, 20 mM imidazole and 0.1% w/v DDM/0.01% w/v CHS), once with 40 mL of Buffer C (50 mM HEPES pH 7.5, 300 mM NaCl, 5% glycerol, 1 mM TCEP, 75 mM imidazole and 0.1% w/v DDM/0.01% w/v CHS) and eluted with 4 mL of buffer D (50 mM HEPES pH 7.5, 300 mM NaCl, 5% glycerol, 1 mM TCEP, 260 mM imidazole and 0.1% w/v DDM/0.01% w/v CHS). Protein concentration was verified using the NanoDrop100 Spectrophotometer (Thermo Fisher Scientific).

### Construction of NahK recombinant plasmids

The gene encoding the C-terminal 6xHis GlcNAc-1-kinase NahK from *Bifodobacterium longum* JCM1217 was inserted into a pBAD/HisB backbone using a NEBuilder^®^ HiFi DNA Assembly Cloning Kit (New England Biolabs) following manufacturer’s protocol. Briefly, using Q5 DNA polymerase (New England Biolabs) the *nahK* gene to be inserted was amplified from plasmid pET22b-6xHis-*nahK*, and the pBAD destination vector was amplified from plasmid pBAD-*gaf2* following manufacturer’s protocols. PCR reactions were cleaned up using a QIAquick PCR Purification Kit (QIAGEN) and ligated with the NEBuilder^®^ HiFi DNA Assembly Cloning Kit using a 1:2 (mol/mol) ratio of vector/insert. Reaction mixture (0.5 µL) was used to transform MAX Efficiency™ DH5 Competent Cells (Invitrogen), and positive transformants were identified by colony PCR using FroggaBio Taq DNA polymerase (FroggaBio). Plasmid sequence was confirmed by Sanger sequencing, performed at the Centre for Applied Genomics at SickKids Hospital for Sick Children.

To generate the inactive NahK^D208N/K210E/N213A^ triple mutant, a Q5 Site-Directed Mutagenesis Kit was used, with all 3 point mutations encoded by either the forward or reverse mutagenic primers.

All recombinant plasmids were transformed into appropriate cell strains by calcium chloride heat shock transformation.

### PgaCD dot blot assay

Reactions contained purified PgaCD_m_ (20 mg/L), MgCl_2_ (5 mM), UDP-GlcNAc (0.3 mM), and C6-substituted UDP-GlcNAc derivative (0-10 mM) in buffer (50 mM HEPES at pH 7.5, 0.3 M NaCl, and 5% glycerol). The reactions were initiated by the addition of substrate. After incubation for 1 h at rt aliquots (5 μL) were blotted onto nitrocellulose. The blots were air-dried (1 h), blocked with 5% BSA in PBS-T (0.1% Tween 20, 15 mL, 1 h, rt), and treated with HRP-WGA (0.5 mg/L, 1 h, rt). The blot was washed three times (15 mL PBS-T, 10 min, rt) and WGA was visualized using SuperSignal™ West Pico PLUS Chemiluminescent Substrate (Thermo Scientific).

Densitometry was used to convert the blot images into dose-response curves. Each spot was integrated in Gel Analyzer. A low level of background signal resulted from the enzyme preparation and was subtracted from each data point before analysis. The data was fit to the IC_50_ equation:

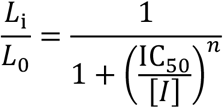

Where *L*_i_/*L*_0_ is the ratio of luminescence of the inhibited and control reactions, [I] is the inhibitor concentration, and *n* is the hill co-efficient. All nonlinear regression analysis was done using the program *KaleidaGraph* v. 4.03 from Synergy Software (Reading, PA).

### PgaCD inhibition time course

Reactions (200 μL) contained PgaCD_m_(MP) (80 mg/mL), MgCl_2_ (5 mM), UDP-GlcNAc (10 mM), and UDP-6-deoxyGlcNAc (0-75 μM) in buffer (50 mM HEPES at pH 7.5, 0.3 M NaCl, 1 mM TCEP and 5% glycerol). The reactions were initiated by the addition of substrate and incubated at 37 °C. Aliquots (20 μL) were taken at 0, 1, 2, 4, 6, 8 and 22 h and stored at −20 °C until analysis. The samples were boiled (5 min), clarified (16,000 x *g*, 10 min), and blotted (5 μL) onto nitrocellulose and analyzed as for the PgaCD dot blot assay.

### PgaCD dialysis experiment

Reactions (200 μL) contained PgaCD_m_(MP) (85 mg/mL), MgCl_2_ (5 mM), UDP-GlcNAc (0 or 10 mM), and UDP-6-deoxyGlcNAc derivative (0 or 0.5 mM) in buffer (50 mM HEPES at pH 7.5, 0.3 M NaCl, 1 mM TCEP and 5% glycerol). The reactions were initiated by the addition of substrate and incubated at 37 °C for 17 h. The samples were dialyzed (25 kDa cut-off) against buffer (1 L, 4 °C) for 8 h. The buffer was replaced (1 L, 4 °C) and the samples dialyzed for a further 16 h. The recovered samples were supplemented with MgCl_2_ (5 mM) and UDP-GlcNAc (10 mM). They were incubated at 37 °C and aliquots (20 μL) were taken at 0, 1, 2, 4, 7, and 23 h, storing them at −20 °C until analysis. The samples were boiled (5 min), clarified (16,000 x *g*, 10 min), and blotted (5 μL) onto nitrocellulose and analyzed as for the PgaCD dot blot assay.

### PgaCD dilution experiment

Reactions (200 μL) contained PgaCD_m_(MP) (85 mg/mL), MgCl_2_ (5 mM), UDP-GlcNAc (0 or 2 mM), and UDP-6-deoxyGlcNAc derivative (0 or 50 μM) in buffer (50 mM HEPES at pH 7.5, 0.3 M NaCl, 1 mM TCEP and 5% glycerol). The reactions were initiated by the addition of substrate and incubated at 37 °C for 17 h. The samples were diluted 10-fold into fresh buffer supplemented with MgCl_2_ (5 mM) and UDP-GlcNAc (10 mM) and incubated at 37 °C. Aliquots (20 μL) were taken at 0, 1, 2, 4, 7, and 23 h and stored at −20 °C until analysis. The samples were boiled (5 min), clarified (16,000 x *g*, 10 min), and blotted (5 μL) onto nitrocellulose and analyzed as for the PgaCD dot blot assay.

### Disaccharide product detection by ESI-HRMS

Reactions (200 μL) contained PgaCD_m_(MP) (72 mg/mL), MgCl_2_ (5 mM), GlcNAc (50 mM), and C6-substituted UDP-GlcNAc analogue (5 mM) in buffer (50 mM HEPES at pH 7.5, 0.3 M NaCl, 1 mM TCEP and 5% glycerol). The samples were incubated at 37 °C for 48 h then boiled (5 min), clarified (16,000 x *g*, 10 min), and loaded onto a pre-equilibrated Grace Alltech Extract-Clean Carbograph column. The column was washed with H_2_O (6 mL) and PNAG oligomers were eluted with 25% v/v acetonitrile (6 mL).^78^ The samples were lyophilized and subjected to ESI-HRMS analysis.

### Biofilm assay

An overnight culture of *E. coli* MG1655 *csrA::kanB* pBAD-*nahK* grown with ampicillin (100 mg/L) and kanamycin (50 mg/L) was diluted to OD_600_ = 0.02 in fresh LB media with ampicillin (100 mg/L), kanamycin (50 mg/L), arabinose (0-200 mg/L), and GlcNAc derivative (0-5 mM). Biofilms were grown in the wells of a 96-well plate (200 μL) for 24 h at rt without shaking. The media was shaken out of the wells and the plate was washed once with distilled water. Adherent biomass was stained with crystal violet (0.1% w/v, 200 μL, rt, 30 min). The stain was shaken out of the wells and the plate washed once with distilled water. The wells were air-dried (4-16 h) and the crystal violet was solubilized in 30% acetic acid (200 μL, rt, 30 min). The resulting solutions were diluted (30-50 μL into 200 μL of 30% acetic acid) and the absorbance at 550 nm measured by plate reader. Each experimental condition was replicated 3-6 times on the same plate. The relative biomass was calculated by taking the ratio of the average crystal violet signal under the experimental condition to the positive control. Biofilm assays with *E. coli* MG1655 *csrA::kanB* without pBAD vector were conducted as above but without ampicillin.

Since the biofilm inhibition plateaued at high concentrations, the dose-response curves were fit using the EC_50_ model:

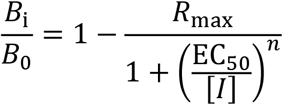

Where *B*_i_/*B*_0_ is the ratio of biofilm biomass with and without the inhibitor, *R*_max_ is the maximum fractional reduction in biomass, EC_50_ is the inhibitor concentration required to achieve 50% maximum effect, [I] is the inhibitor concentration, and *n* is the Hill coefficient. The maximum level of biofilm inhibition observed varied and could not always be accurately measured. Therefore, we reported IC_50_ values based on the EC_50_ curve fit to allow the relative activities of the inhibitors to be fairly compared:

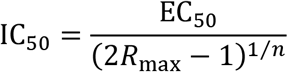

### Growth curves

*E. coli* MG1655 *csrA::kanB* pBAD-*nahK* was grown at 37 °C with shaking (200 rpm) in LB media (2 mL) with ampicillin (100 mg/L), kanamycin (50 mg/L) and L-arabinose (50 mg/L) from an initial OD_600_ of 0.02. Before each OD_600_ measurement the culture was mixed by pipetting to ensure homogeneity. Above OD_600_ ∼ 0.7 a 1/10 dilution of the culture into fresh LB media was measured.

### ^1^H NMR NahK activity assay

Reactions contained GlcNAc derivative (10 mM), ATP (10 mM), MgCl_2_ (5 mM), Tris (0.1 M, pH 8), and a variable amount of NahK (1, 5 or 25 μM). The reactions were heated to 37 °C and allowed to proceed for 1 h or 20 h. The reactions were stopped by boiling (1 min) followed by centrifugation (16,000 x *g*, 10 min). The clarified supernatant was lyophilized and dissolved in D_2_O (150 μL). ^1^H NMR spectra (64 scans, 1 s relaxation delay) were recorded on a 400 MHz Bruker Advance III spectrometer. Reaction progress was estimated by taking the integration ratio of the GlcNAc 1-phosphate H-1 peak (∼5.4 ppm) and the adenosine ribose H-1 signal (6.15 ppm).

### In vivo PNAG extraction/quantitation

Cultures of *E. coli* were grown in LB media (1 mL) with ampicillin (100 mg/L), kanamycin (50 mg/L), arabinose (0-50 mg/L) and 6-fluoroGlcNAc (0-200 μM). Beginning from an initial OD_600_ = 0.02 the strains were grown under static, aerobic conditions at rt for 24 h. The cultures were spun down (6000 x *g*, 3 min). The cell pellets were resuspended in EDTA (50 μL, 0.5 M, pH 8) and boiled (5 min). The samples were clarified (16,000 x *g*, 5 min) and the supernatant (40 μL) was treated with proteinase K (10 μL, 20 mg/mL, 37 °C, 0.5 h). These samples were blotted (5 μL) onto nitrocellulose and detected as for the PgaCD dot blot assay.

### GFP-DspB^*E184Q*^ *PNAG binding assay*

Overnight cultures of *E. coli* MG1655 *csrA::kanB* pBAD-*nahK/nahK*_m_ grown with ampicillin (100 mg/L) and kanamycin (50 mg/L) were used to inoculate fresh LB media (5 mL) with ampicillin (100 mg/L), kanamycin (50 mg/L), arabinose (50 mg/L), and 6-fluoroGlcNAc (0-200 μM). Cultures were grown at 37 °C with shaking to a final OD_600_ of ∼1, harvested (6500 x *g*, 30 min, 4 °C) and washed once with PBS. Cells were resuspended in assay buffer (50 mM sodium phosphate pH 5.8, 0.5% (w/v) BSA, PBS pH 5.8) to an OD_600_ = 5.0. Cell samples were applied (0.5 mL) to microcentrifuge 0.45 μm spin filter tubes (ref. CLS8163, Corning Costar); all centrifugation steps with filter tubes were performed at 4 °C and 4000 x *g*. Samples were filtered by centrifugation, resuspended in assay buffer with GFP-DspB^E184Q^ (0.1 μM, 150 μL), and incubated to label PNAG (rt, 10 min). Samples were again filtered by centrifugation, resuspended in assay buffer (150 μL), and incubated to wash (rt, 10 min). In the final step, samples were filtered by centrifugation, resuspended in assay buffer with DspB enzyme (50 μM, 150 μL), and incubated to hydrolyze PNAG (rt, 60 min). After filtration by centrifugation, GFP fluorescence (λ_ex_: 470 nm, λ_em_: 515 nm) of the collected filtrates were measured using a CLARIOstar microplate reader (BMG Labtech). Values were background corrected using an *E. coli* MG1655 *csrA::kanB* pBAD-*nahK* cell sample treated by a modified protocol without added GFP-DspB^E184Q^, and normalized to the initial labeling concentration of GFP-DspB^E184Q^ (0.1 μM). Each experimental condition was repeated once per biological replicate, with 4 biological replicates performed.

### E. coli metabolite extract and clean up

*E. coli* MG1655 *csrA::kanB* pBAD-*nahK* was grown at 37 °C with shaking (200 rpm) in LB media (20 mL) with ampicillin (100 mg/L), kanamycin (50 mg/L), L-arabinose (50 mg/L), and 6-fluoroGlcNAc (0, 20, or 200 μM) from initial OD_600_ of 0.02. At OD_600_ = 1.0, the cells were harvested (6000 x *g*, 3 min, 4 °C) and washed once with PBS (1 mL, 4 °C). The cell pellets were stored at −20 °C until extraction.

The cell pellets were thawed and suspended in 5% TCA (100 μL, rt, 15 min). Next, the samples were clarified (16,000 x *g*, 5 min, 4 °C). The supernatant was diluted into potassium phosphate (1 mL, 10 mM, pH 2.5) and the pH adjusted to within 2.5-4.5 by addition of KOH (20 to 30 μL, 1 M). The samples were applied to 3 mL Supelco SPE-SAX columns. Before use, the columns were conditioned with MeOH (2 mL), water (2 mL), and 10 mM potassium phosphate (2 mL, pH 2.5). The columns were washed with 10 mM potassium phosphate (5 mL, pH 2.5), 50 mM (2.5 mL, pH 2.5), then 150 mM (0.75 mL, pH 7.5). The UDP-sugars were eluted with another portion of 150 mM potassium phosphate (1 mL, pH 7.5). The samples were stored at −80 °C until HPLC analysis.

### HPLC method

A Restek Ultra IBD column (5 µm, 250 × 4.6 mm) was ran isocratically with mobile phase: 17.5 mM potassium phosphate, 2 mM Bu_4_N-phosphate, pH 6.2. The column temperature was 25 °C, the flow rate was 1 mL/min, the injection volume was 10 µL, and detection was done by absorbance at 262 nm. Before each lysate run the column was first equilibrated with mobile phase (3 h). Next, a series of standards were run including UDP-GlcNAc (8, 4, 2, and 1 µM) and UDP-6-fluoroGlcNAc (10 µM). After analyzing a lysate sample the column was washed as follows: 100% H_2_O, 20 min; 0 to 100% MeOH gradient, 40 min; 100% MeOH, 30 min. The column was stored in 50% MeOH.

### Chemical synthesis

The synthesis of the C6-substituted GlcNAc derivatives was previously reported,^56^ except for 6-aminoGlcNAc which is described below.

#### 2-Acetamido-6-amino-,6-dideoxy-D-glucopyranoside (1-NH_2_)

2-Acetamido-6-azido-2,6-dideoxy-D-glucopyranoside (6.0 mg, 0.024 mmol) and Pd/C (5 mg, 10%) were suspended in water (1 mL) acidified with TFA (2 eq., 48 mM). The reaction was purged with N_2_(g) followed by H_2_(g) and stirred at rt for 2 h. The mixture was filtered through celite and lyophilized, giving the product as its TFA salt (7.6 mg, 93%, white solid). 1H NMR (400 MHz, D_2_O) δ 5.22 (d, *J* = 3.5 Hz, 0.61H, αH-1), 4.75 (d, *J* = 8.6 Hz, 0.41H, βH-1), 4.04 (td, *J* = 9.4, 3.1 Hz, 0.63H), 3.98 – 3.83 (m, 0.81H), 3.81 – 3.62 (m, 1.35H), 3.56 (dd, *J* = 10.4, 8.8 Hz, 0.34H), 3.52 – 3.33 (m, 1.96H), 3.16 (ddd, *J* = 13.2, 8.9, 4.1 Hz, 0.89H), 2.05 (s, 3H). ^13^C NMR (126 MHz, D_2_O) δ 174.7, 174.5, 162.9 (q, *J* = 35.7 Hz, TFA), 116.3 (q, *J* = 291.7 Hz, TFA), 94.8 (βC-1), 90.8 (αC-1), 73.4, 71.8, 71.6, 71.4, 70.3, 67.3, 56.4, 53.8, 40.4, 40.3, 22.1, 21.8. HRMS(ESI) *m/z* Calcd. for C_8_H_17_N_2_O_5_ [M+H]^+^: 221.1132, found 221.1129.

## Author contributions

Z.A.M., A.E., M.N. conceptualization; Z.A.M, A.E., A.S.S, investigation; Z.A.M., A.E., M.N. writing-original draft; Z.A.M, A.E., A.S.S, and M.N.

P.L.H. writing-review and editing; M.N. and P. L. H. supervision; M.N. and P. L. H. funding acquisition; M. N. project administration.

## Funding and additional information

This work was supported in part by Natural Sciences and Engineering Research Council (NSERC) Grant (to M.N.). Canadian Institutes of Health Research (CIHR) Grants FDN154327 (to P. L. H.), P. L. H. is a recipients of Canada Research Chair (2006-2020). This work has also been supported by Canada Graduate Scholarships from NSERC (to A.S.S and Z.A.M.), the Ontario Graduate Scholarship Program (A.E.) The Hospital for Sick Children Foundation Student Scholarship Program (to A.S.S),

