## Supporting Information for "Truncation of poly-*N*-acetylglucosamine (PNAG) polymerization with *N*-acetylglucosamine analogues"

### Table of Contents


| | |
|---|---|
| Figure S1 | Purification of PgaCDm and validation of dot blot assay |
| Figure S2 | Representative dot blot dose-response assays for inhibition of purified PgaCDm by C6-substituted UDP-GlcNAc derivatives |
| Figure S3 | L-Arabinose-induced expression of (A) pBAD-nahK and (B) pBAD-nahKm in *E. coli csrA::kanB* |
| Figure S4 | NahK activity does not perturb biofilm formation or growth. |
| Figure S5 | Growth curves of *E. coli csrA::kanB pBAD-nahK* in the presence of C6-substituted GlcNAc derivatives |
| Figure S6 | Confirmation of HPLC peak identities |
| Figure S7 | Metabolite extract chromatograms for *E. coli* MG1655 csrA::kanB pBAD-nahK |
| Table S1 | Plasmid and strain table |

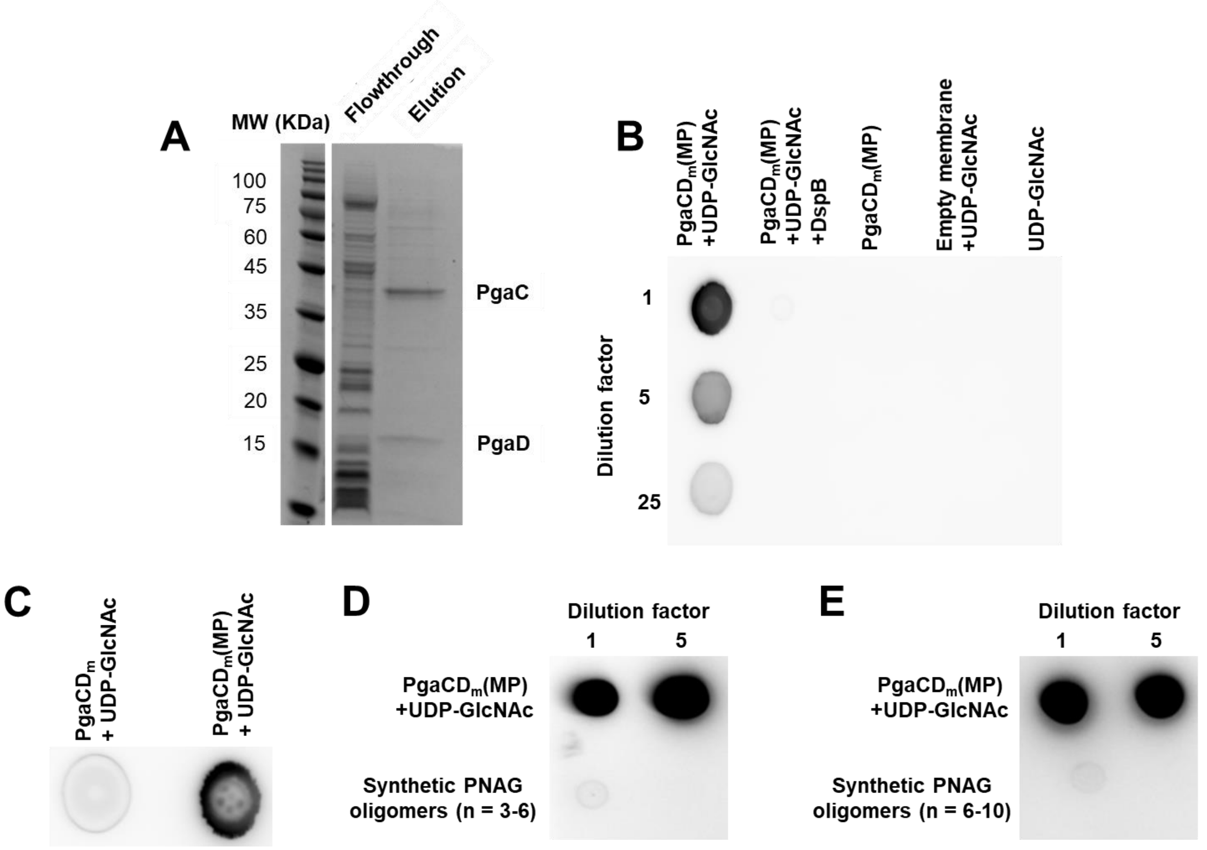


**Figure S1. Purification of PgaCD$_m$ and validation of dot blot assay. A**) Nickel affinity purification of PgaCD$_m$. The bands at ~45 kDa and ~16 kDa correspond to PgaC and PgaD respectively. **B**) Dot blot assay HRP signal requires PgaCD$_m$ activity and UDP-GlcNAc. DspB treatment of the PgaCD$_m$(MP)/UDP-GlcNAc reaction abolishes the HRP signal. DspB reaction conditions: DspB (5 µM), $NaHPO_4$ (100 mM), pH 5.8, 2 h, 37 °C. **C**) *E. coli* membrane preparation overexpressing PgaCD$_m$ (PgaCD$_m$(MP)) has greater activity than purified PgaCD$_m$. Conditions: membrane preparation (90 mg/mL), or purified PgaCD$_m$ (0.25 mg/mL) UDP-GlcNAc (10 mM), $MgCl_2$ (5 mM), 37 °C, 24 h. **D-E**) Synthetic PNAG oligomers (8.7 mg/mL, 5 µL) of lengths n = 3-6 (**D**) or n = 6-10 (**E**), show minimal WGA-HRP signal compared to PgaCD membrane preparation/UDP-GlcNAc reaction.

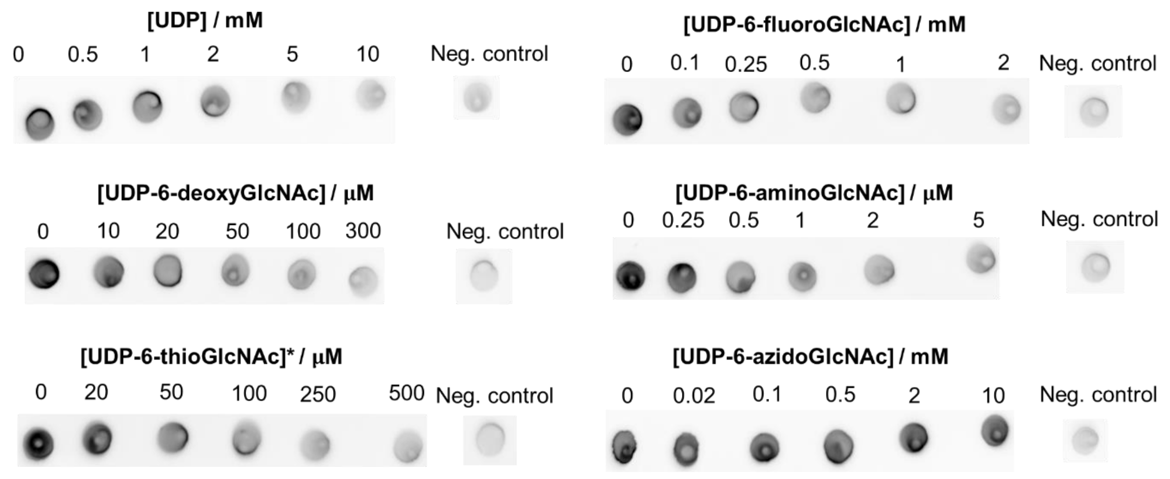


**Figure S2**. **Representative dot blot dose-response assays for inhibition of purified PgaCD$_m$ by C6-substituted UDP-GlcNAc derivatives.** The negative controls are reaction mixture without substrate. Conditions: UDP-GlcNAc (0.3 mM), PgaCD (20 mg/L), $MgCl_2$ (5 mM), rt, 1 h. *Assayed in the presence of 5 mM DTT to maintain a reducing environment.

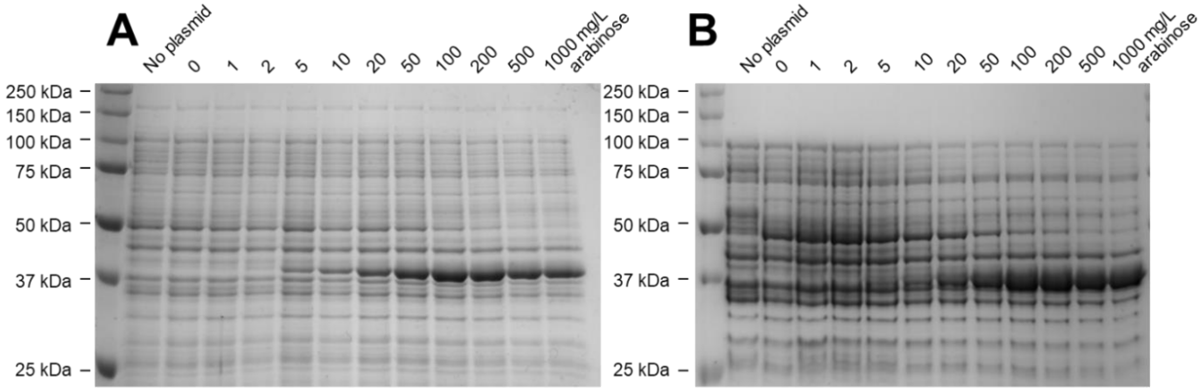


**Figure S3**. **L-Arabinose-induced expression of (A) pBAD-*nahK* and (B) pBAD-*nahK*$_m$ in *E. coli csrA*::*kanB*.** Whole cleared cell lysates were analyzed directly by SDS-PAGE gel. NahK runs as a band at 41 kDa. Biological replicates of both experiments gave qualitatively identical results.

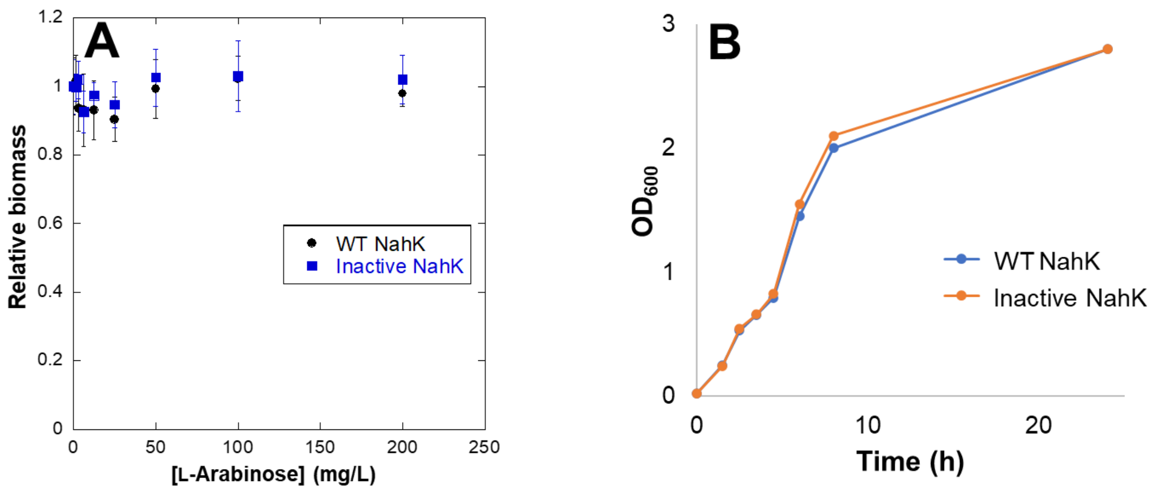


**Figure S4**. **NahK activity does not perturb biofilm formation or growth. A**) Induction of *E. coli csrA*::*kanB* pBAD-*nahK* or *E. coli csrA*::*kanB* pBAD-*nahK*m*)* with L-arabinose does not appreciably affect biofilm formation. **B**) pBAD-*nahK* or pBAD-*nahK*m gives equivalent growth rates in *E. coli csrA*::*kanB* with 50 mg/L L-arabinose.

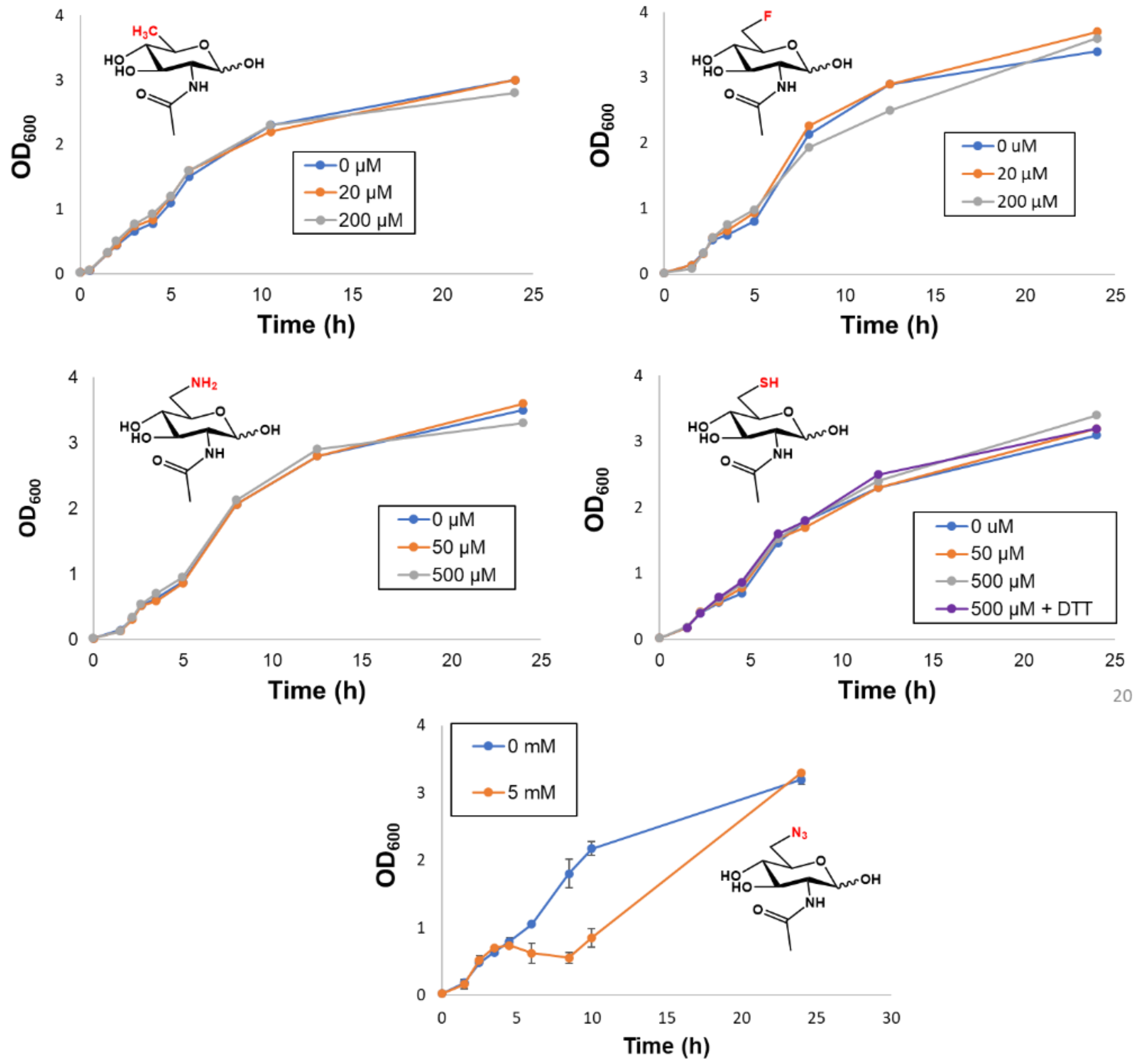


**Figure S5**. **Growth curves of *E. coli csrA*::*kanB* pBAD-*nahK* in the presence of C6-substituted GlcNAc derivatives.** NahK was induced at 50 mg/L L-arabinose. The 6-azidoGlcNAc growth curve is the average of two biological replicates.

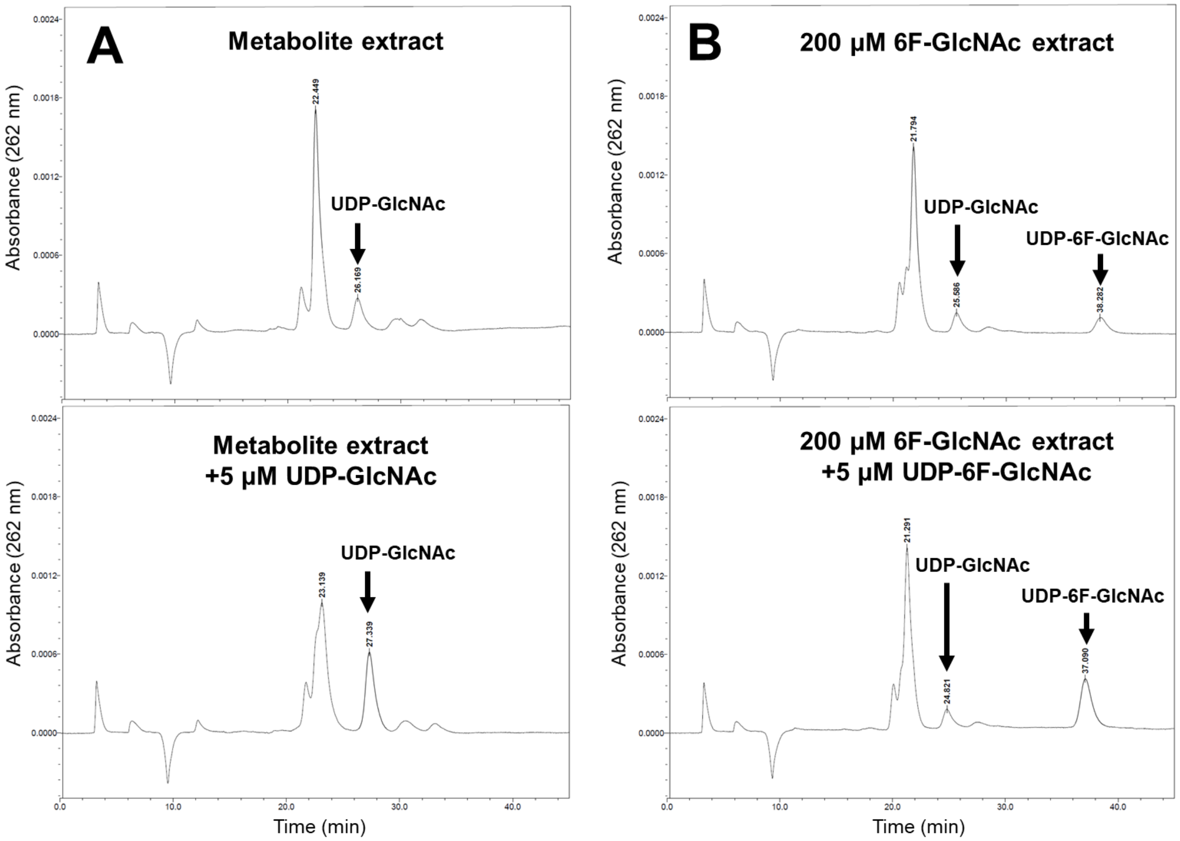


**Figure S6. Confirmation of HPLC peak identities.** Chromatograms of *E. coli* MG1655 *csrA::kanB* pBAD-*nahK* metabolite extracts before and after standard addition to confirm the identity of the UDP-GlcNAc (**A**) and UDP-6-fluoroGlcNAc (**B**) peaks.

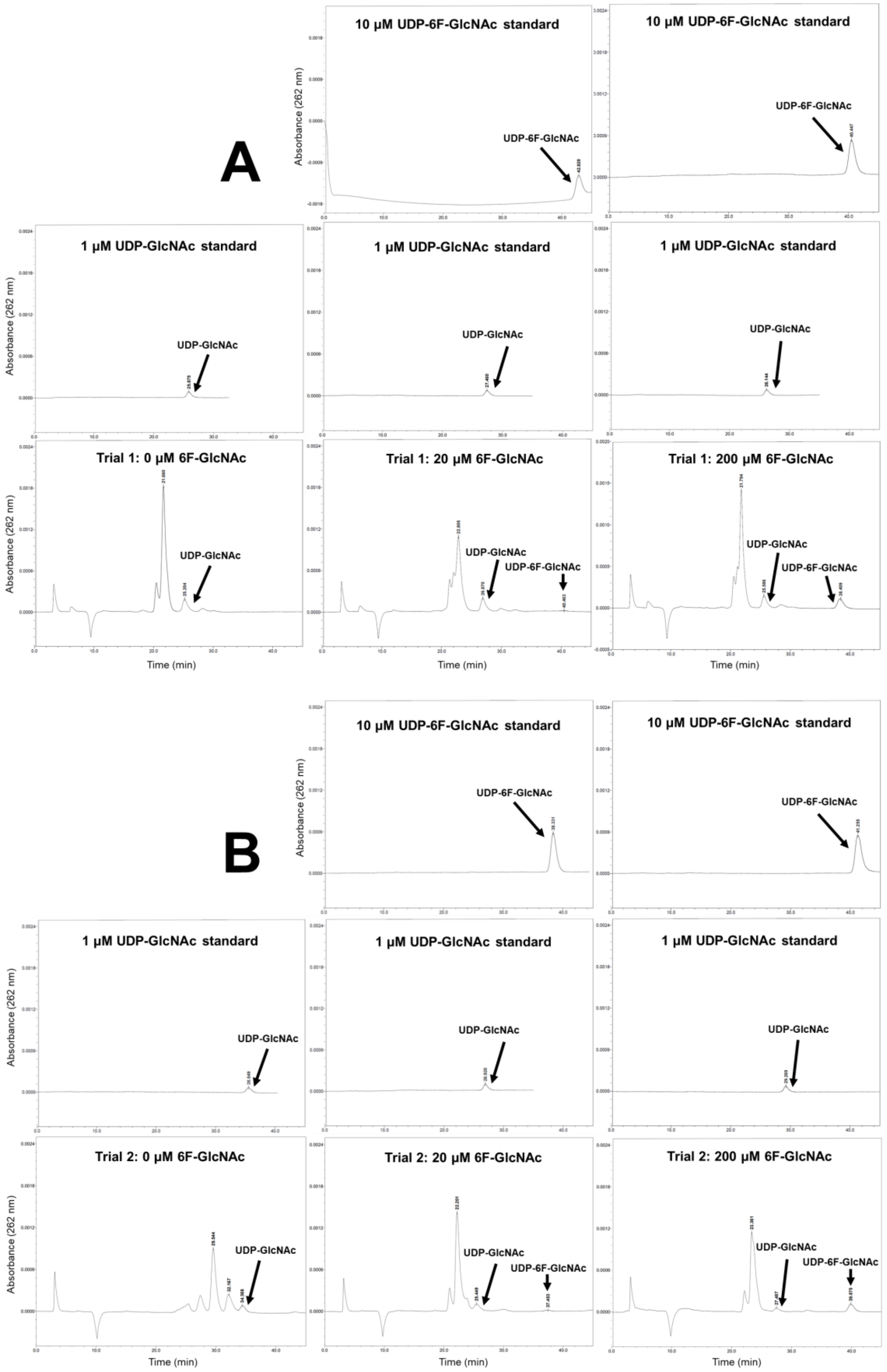
A
10 µM UDP-6F-GlcNAc standard
10 µM UDP-6F-GlcNAc standard
UDP-6F-GlcNAc
UDP-6F-GlcNAc
1 µM UDP-GlcNAc standard
1 µM UDP-GlcNAc standard
1 µM UDP-GlcNAc standard
UDP-GlcNAc
UDP-GlcNAc
UDP-GlcNAc
Trial 1: 0 µM 6F-GlcNAc
Trial 1: 20 µM 6F-GlcNAc
Trial 1: 200 µM 6F-GlcNAc
UDP-GlcNAc
UDP-GlcNAc
UDP-6F-GlcNAc
UDP-GlcNAc
UDP-6F-GlcNAc
Absorbance (262 nm)
Time (min)
B
10 µM UDP-6F-GlcNAc standard
10 µM UDP-6F-GlcNAc standard
UDP-6F-GlcNAc
UDP-6F-GlcNAc
1 µM UDP-GlcNAc standard
1 µM UDP-GlcNAc standard
1 µM UDP-GlcNAc standard
UDP-GlcNAc
UDP-GlcNAc
UDP-GlcNAc
Trial 2: 0 µM 6F-GlcNAc
Trial 2: 20 µM 6F-GlcNAc
Trial 2: 200 µM 6F-GlcNAc
UDP-GlcNAc
UDP-GlcNAc
UDP-6F-GlcNAc
UDP-GlcNAc
UDP-6F-GlcNAc
Absorbance (262 nm)
Time (min)

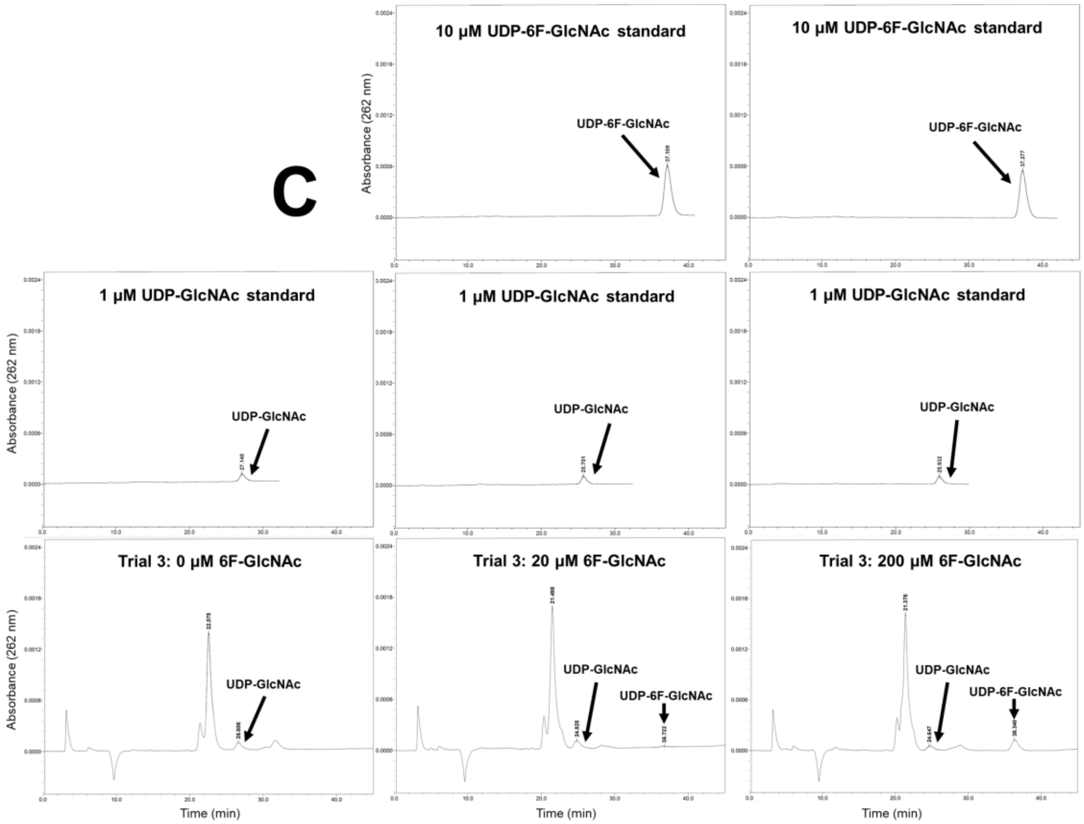


**Figure S7**. **Metabolite extract chromatograms for *E. coli* MG1655 *csrA::kanB* pBAD-*nahK.*** Three biological replicates (**A**-**C**) were performed. The cells were grown with 0, 20, or 200 µM 6-fluoroGlcNAc and L-arabinose (50 mg/L). Peaks were assigned using the retention times of standards ran immediately before the lysate samples, provided above each lysate chromatogram.

| Plasmid or Strain | Description | Origin |
|---|---|---|
| **Plasmid** | | |
| pBAD-*nahK* | Arabinose-inducible expression plasmid for NahK | This paper |
| pBAD-*nahK*$_m$ (D208N/K210E/N213A triple mutant) | Arabinose-inducible expression plasmid for NahK$_m$ | This paper |
| pET29b-*dspB* | IPTG-inducible expression plasmid for DspB | Narayanan Ramasubbu lab, Department of Oral Biology, University of Medicine and Dentistry of New Jersey |
| pET29b-*egfp-dspB*$^{E184Q}$ | IPTG-inducible expression plasmid for EGFP-DspB$^{E184Q}$ | Eddenden et al., 2020 (Addgene#175802) |
| pET22b-*nahK* | IPTG-inducible expression plasmid for NahK | Peng George Wang lab, Departments of Chemistry and Biochemistry, The Ohio State University |
| pET22b-*nahK*$_m$ (*nahK*$^{D208N/K210E/N213A}$) | IPTG-inducible expression plasmid for NahK$_m$ | This paper |
| pET15b-*glmU* | IPTG-inducible expression plasmid for GlmU | Peng George Wang lab, Departments of Chemistry and Biochemistry, The Ohio State University |
| pBAD-*gaf2* | Backbone used in Gibson assembly to generate pBAD-*nahK* | Andrew Woolley lab, Department of Chemistry, University of Toronto |
| pQLinkH-6His-*pgaC* | IPTG-inducible expression plasmid for PgaC | This paper |
| pQLinkN-*pgaD* | IPTG-inducible expression plasmid for PgaD | This paper |
| pQLinkN-*pgaD*-FLAG | IPTG-inducible expression plasmid for PgaD-FLAG | This paper |
| pQLink-6His-*pgaC-pgaD*-FLAG | IPTG-inducible co-expression plasmid for PgaCD | This paper |
| **Strain** | | |
| Escherichia coli NEB5α | *fhuA2 Δ(argF-lacZ)U169 phoA glnV44 Φ80 Δ(lacZ)M15 gyrA96 recA1 relA1 endA1 thi-1 hsdR17* | New England Biolabs (Cat#2987H) |
| *Escherichia coli* BL21(DE3) | *fhuA2 [lon] ompT gal (λ DE3) [dcm] ∆hsdS λ DE3 = λ sBamHIo ∆EcoRI-B int::(lacI::PlacUV5::T7 gene1) i21 ∆nin5* | New England Biolabs (Cat#2527H) |
| *Escherichia coli* MG1655 *csrA*::*kanB* | *K-12 F– λ– ilvG– rfb-50 rph-1 csrA*::*kanB* | Tony Romeo lab, Department of Microbiology and Cell Science, University of Florida |

| | | |
|---|---|---|
| *Escherichia coli* BW25113 | *lacI+rrnBT14 ΔlacZWJ16 hsdR514 ΔaraBADAH33 ΔrhaBADLD78 rph-1 Δ(araB–D)567 Δ(rhaD–B)568 ΔlacZ4787(::rrnB-3) hsdR514 rph-1* | Coli Genetic Stock Center (CGSC#7636) |
